# Regulatory annotation identifies KAN1, MYB44 and PIF4 as regulators of Arabidopsis lincRNAs expressed in root

**DOI:** 10.1101/2022.09.09.507345

**Authors:** Li Liu, Michel Heidecker, Thomas Depuydt, Nicolas Manosalva Perez, Martin Crespi, Thomas Blein, Klaas Vandepoele

## Abstract

Thousands of long intergenic noncoding RNAs (lincRNAs) have been identified in plant genomes. While some lincRNAs have been characterized as important regulators in different biological processes, little is known about the transcriptional regulation for most plant lincRNAs. Through the integration of eight annotation resources, we defined 6,599 high-confidence lincRNA loci in *Arabidopsis thaliana*. For lincRNAs belonging to different evolutionary age categories, we identified major differences in sequence and chromatin features, as well as in the level of conservation and purifying selection acting during evolution. Spatiotemporal gene expression profiles combined with transcription factor (TF) chromatin immunoprecipitation data were used to construct a TF- lincRNA regulatory network containing 2,659 lincRNAs and 15,686 interactions. We found that properties characterizing lincRNA expression, conservation and regulation differ between plants and animals. Experimental validation confirmed the role of three TFs, KAN1, MYB44, and PIF4, as key regulators controlling root-specific lincRNA expression, demonstrating the predictive power of our network. Furthermore, we identified 58 lincRNAs, regulated by these TFs, showing strong root cell-type specific expression or chromatin accessibility, which are linked with GWAS genetic associations related to root system development and growth. The multi-level genome-wide characterization covering chromatin state information, promoter conservation, and ChIP-based TF binding, for all detectable lincRNAs across 769 expression samples, permits to rapidly define the biological context and relevance of lincRNAs in Arabidopsis regulatory networks.

**One-line sentence:** A multi-level Arabidopsis gene regulatory network identifies novel regulators controlling root-specific lincRNA expression, offering a promising strategy to identify lincRNAs involved in plant biology.

## Introduction

Genomes are widely transcribed and produce thousands of long non-coding RNAs (lncRNAs), which are an abundant class of transcripts longer than 200 nucleotides with low protein coding capacity (Wu et al., 2017). LncRNAs are generally transcribed by RNA polymerase II (Pol II), are poly-adenylated and may contain introns as coding RNA. LncRNAs modulate gene expression through a wide range of mechanisms, including chromatin structure remodeling, transcription regulation in cis/trans, fine-tuning of miRNA activity, alternative splicing (AS) regulation, and the control of mRNA stability and translation (Sanchita et al., 2020; Bhogireddy et al., 2021; Lucero et al., 2021). One class of lncRNAs is the primary mRNAs containing miRNA precursors or pri-miRNAs. These are rapidly and generally processed into miRNAs (Jones-Rhoades et al., 2006; Shafiq et al., 2016). However, the large majority of lncRNAs are able to act without being further processed such as the lncRNAs controlling the epigenetic regulation the *Flowering Locus C* (*FLC*) gene expression and mediating plant vernalization, i.e., *COOLAIR*, *COLDAIR* and *COLDWRAP* (Liu et al., 2010; Heo and Sung, 2011; Kim and Sung, 2017). A subgroup of lncRNAs derived from intergenic regions is defined as long intergenic non-coding RNAs (lincRNAs) and have been identified in a wide range of eukaryotes including model and non-model plant species (Wu et al., 2017; Chen et al., 2021). In contrast to antisense lncRNAs, whose sequence evolution is constrained by the overlapping coding genes, the transcription and evolution of lincRNAs are independent of the surrounding genes.

While the low expression levels and tissue-specific expression patterns of lincRNAs in plants initially raised concerns (Liu et al., 2012; Bu et al., 2015), increasing experimental evidence supports the functional activity of lincRNAs. In plants, few lincRNAs have been experimentally validated (Chen et al., 2021), showing their involvement in various biological contexts such as in regulating flowering time (Chen et al., 2020) and root growth and development (Roule et al., 2022a), or influencing germination (Kramer et al., 2022). For example, the Arabidopsis *FLINC* lincRNA has been reported to regulate ambient temperature-mediated flowering time (Severing et al., 2018). Arabidopsis lateral root development is regulated by the *Alternative Splicing COmpetitor* (*ASCO*) lincRNA, which modulates AS by interacting with the multiple splicing factors (Bardou et al., 2014; Rigo et al., 2020), and the *AUXIN-REGULATED PROMOTER LOOP* (*APOLO*) lincRNA, which influences the local chromatin conformation and the activity of several Auxin-Responsive genes (Ariel et al., 2020). Plants lacking *CONSERVED INBRASSICA RAPA1* (*lncCOBRA1*) were found to show a delayed germination and were overall smaller than wild-type plants (Kramer et al., 2022). Many lincRNAs are differentially expressed in various stress responses, including drought (Qi et al., 2013; Shuai et al., 2014; Li et al., 2017; Qin et al., 2017), cold (Li et al., 2017; Zhao et al., 2018; Shea et al., 2019), salinity (Deng et al., 2018; Fukuda et al., 2019), and nutrient deficiency (Fukuda et al., 2019), implying that lincRNAs may be involved in plant stress resilience (Jha et al., 2020). Interestingly, some of the confirmed functional lincRNAs interact with transcription factors (TFs) to activate or repress the expression of associated genes, such as *APOLO* that interacts with WRKY42 to form a regulatory hub that controls the activity of *RHD6* and promotes the expansion of root hair cell at low temperatures (Moison et al., 2021; Pacheco et al., 2021).

Recently, a lot of attention has been placed on the evolutionary conservation of lincRNAs, which is generally associated with functionality (Ulitsky, 2016; Szczesniak et al., 2021). The conservation of noncoding transcripts can be considered at the level of the primary sequence, position, splice sites, RNA structure, and transcriptional level (Ulitsky, 2016; Szczesniak et al., 2021). However, most lincRNA sequences undergo rapid evolution and are poorly conserved (Ransohoff et al., 2018). In a study by Wang et al., only 5% of 2,281 rice lincRNAs had sequence similarity to maize lincRNAs. It was also found that the positional conservation of lincRNAs was much higher than their sequence conservation (Wang et al., 2015). Nelson and co-workers reported that 22% of 1180 Arabidopsis lincRNA loci were conserved in 10 Brassicaceae genomes (Nelson et al., 2016). These conserved lincRNAs exhibited higher expression levels, stress-responsiveness and their gene body overlapped with conserved noncoding sequences (CNSs), suggesting a role of their conserved sequence in a genomic context (Nelson et al., 2016).

While different studies have reported on the identification and expression of lincRNAs in plants (Wang et al., 2015; Nelson et al., 2016; Ke et al., 2019; He et al., 2021), a comprehensive overview of the different genomic features contributing to the expression, regulation and evolutionary conservation of plant lincRNAs is missing. How lincRNAs are embedded in transcriptional networks controlling different biological processes remains largely unknown. Furthermore, prioritizing lincRNAs for downstream functional analysis is not straightforward without knowing the regulatory network where they are involved in. Here, we integrated different Arabidopsis lincRNAs gene annotations and explored various functional genomics datasets to characterize lincRNA expression in a biologically relevant context. We leveraged large-scale expression datasets and protein-DNA interaction data to study the molecular networks controlling lincRNA gene activity. Combined with evolutionary conservation analysis, we explored the global transcriptional regulatory properties of different evolutionary age categories and, through regulatory network analysis, identified specific TFs controlling lincRNA regulation in roots.

## RESULTS

### Integration of lincRNA annotations in *Arabidopsis*

A substantial number of lncRNA transcripts in *Arabidopsis* have been identified and several publicly available resources for the annotation of lncRNAs have been developed (Jha et al., 2020). In contrast to antisense lncRNAs which generally regulate the overlapping coding gene, much less is known about the potential targets of lincRNAs, so we focus our study on these transcripts. Indeed, lincRNAs transcription and evolution are independent of the surrounding genes, in contrast to antisense lncRNAs that are constrained by the coding genes they overlap with. To integrate and unify previously identified lincRNAs, annotations based on transcriptome information from 10 different tissues and various environmental conditions were collected from different resources including Araport11 (Cheng et al., 2017), CANTATAdb (Szczesniak et al., 2016), NONCODEv5 (Fang et al., 2018), PLNlncRbase (Xuan et al., 2015), key lncRNA research articles (Liu et al., 2012; Nelson et al., 2017; Zhao et al., 2018), and new predictions based on root related stranded RNA-seq (Material and Methods). Next, a pipeline was designed to define a unified set of high-confidence lincRNAs by discarding transcripts with length below 200 bp, removing transcripts that overlapped with protein-coding genes (antisense lncRNAs), re-evaluating the coding potential of the transcripts, and merging the remaining transcripts from various resources (see Materials and Methods, Supplemental Figure S1A-B). In total, we identified 7,706 lincRNA transcripts covering 6,599 high-confidence lincRNA loci (see Supplemental Data Set S1). To explore the overlap of this high-confidence lincRNA gene set with the individual annotations, we assessed the overlap between the different resources (Figure 1A). A total of 4,955 (75.1%) lincRNA loci were supported by only one resource and the remaining 1,644 (24.9%) lincRNA loci were derived from two or more resources (Figure 1B). Araport11 contained the highest number of shared loci and Liu et al. (2012) contained the highest number of unique loci (Chen et al., 2021). Next, the genomic features of the high-confidence lincRNA transcripts were compared to those of protein-coding transcripts. 6,428 (83%) lincRNA transcripts and 6,250 (13%) protein-coding transcripts contained single exons, while 1,278 (17%) lincRNA transcripts and 42,109 (87%) protein-coding transcripts contained multiple exons. Furthermore, a higher frequency of multi-exon transcripts was found in lincRNAs supported by two or more resources (974, 36%) than in those supported by a unique resource (304, 6%) (Figure 1C). The transcript length distribution for lincRNAs showed a U-shape curve with the majority of transcripts being 200-300bp long (Figure 1D).

**Figure 1.**
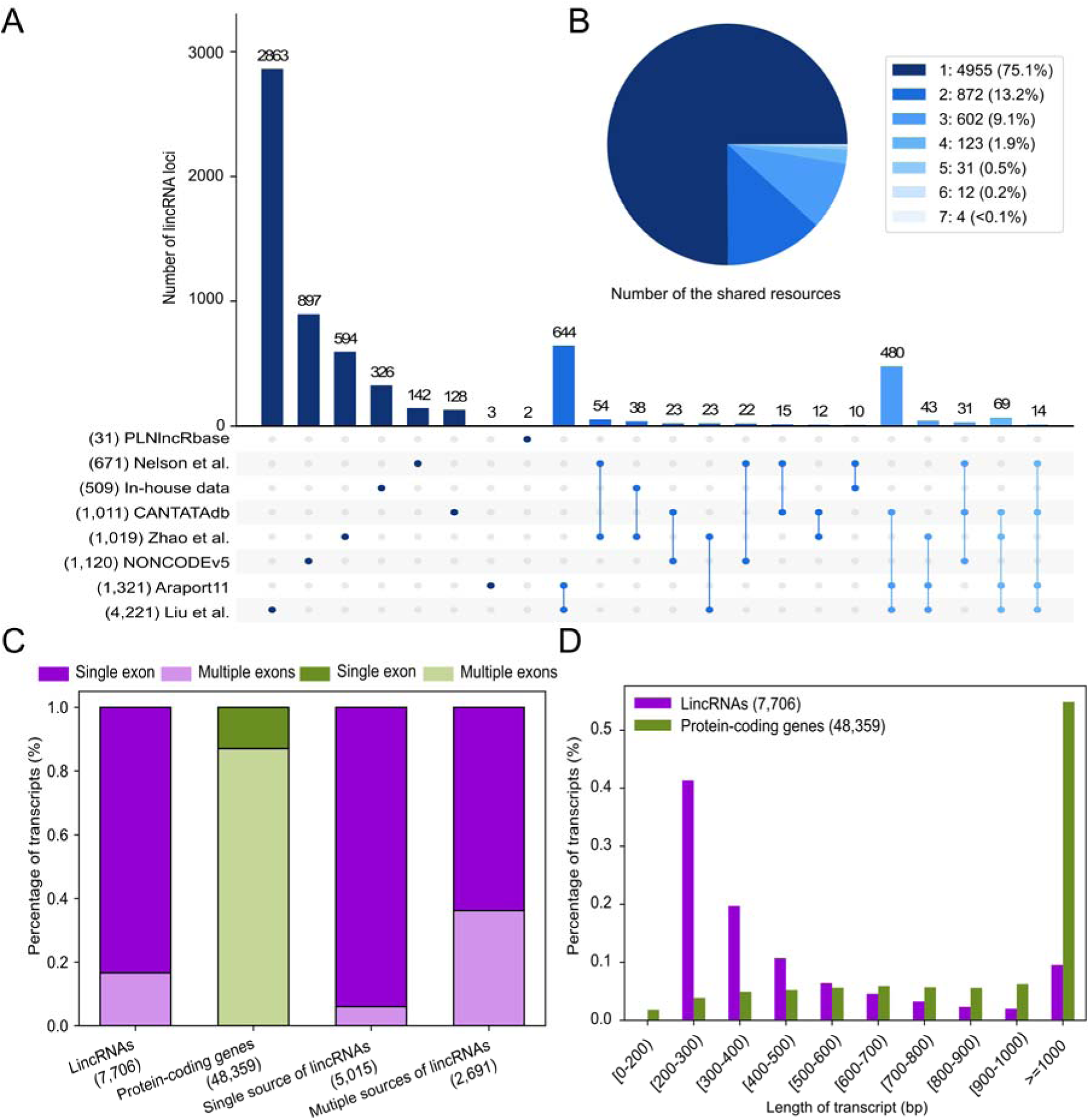
Overlap and gene features of Arabidopsis lincRNA annotations. **(A)** Upset plot showing the intersection of lincRNA annotation in the eight resources. Each row represents a resource, reporting in parenthesis its total number of lincRNA transcripts before merging. LincRNA annotations unique to a single resource are represented as a single circle while circles connected by lines represent the intersection of lincRNA loci shared between various resources. The bar chart indicates the number of unique lincRNA loci and intersectional lincRNA loci, displaying only intersections that contain at least ten lincRNA loci. More complex overlapping patterns are not shown. **(B)** The pie chart shows the proportion of lincRNA loci supported by one or more resources. **(C)** The distribution of exon number for all lincRNA transcripts (purple), protein-coding transcripts (green), transcripts of lincRNAs supported by single resource (purple) and multiple resources (purple). Single exon and multiple exons are shown in dark and light colors, respectively. **(D)** The distribution of transcript length for lincRNAs (purple) and protein-coding genes (green).

### Contrasting patterns of sequence conservation for lincRNAs belonging to different evolutionary age categories

To assess the evolutionary conservation of Arabidopsis lincRNAs within flowering plants, DNA sequence similarity searches were performed by comparing our set of lincRNAs with the genomes of 40 plant species (see Materials and Methods, Supplemental Table S1). Among the 6,599 lincRNAs, 2,480 lincRNAs were restricted to Arabidopsis and named Arabidopsis-specific lincRNAs. The other lincRNAs were classified into four evolutionary age categories according to the presence of homologs in closely and more distantly related species (Figure 2A). We found 81 lincRNAs with at least one homolog in eudicots and in monocots and therefore conserved during 180 million years (MY) of evolution (Beilstein et al., 2010; Zhang et al., 2020), defined as angiosperm-conserved lincRNAs. Forty-two lincRNAs were conserved in eudicots with at least one homolog in rosids and asterids, but without homologs in monocots. Similarly, 44 lincRNAs were identified only having homologs in rosids species, outside the Brassicaceae family. As the lincRNAs conserved in eudicots and rosids showed highly similar conservation patterns for gene body and promoter (P<0.376, Mann–Whitney *U* test), we combined these genes in one category, called Eudicot/rosid-conserved lincRNAs (86 genes). 1,671 Brassicaceae_I_II-conserved lincRNAs were present in the common ancestor of Brassicaceae lineages I and II, without homologs outside the Brassicaceae. Lastly, 2,281 lincRNAs were identified only having homologs in the Brassicaceae I lineage (Brassicaceae_I-conserved lincRNAs).

**Figure 2.**
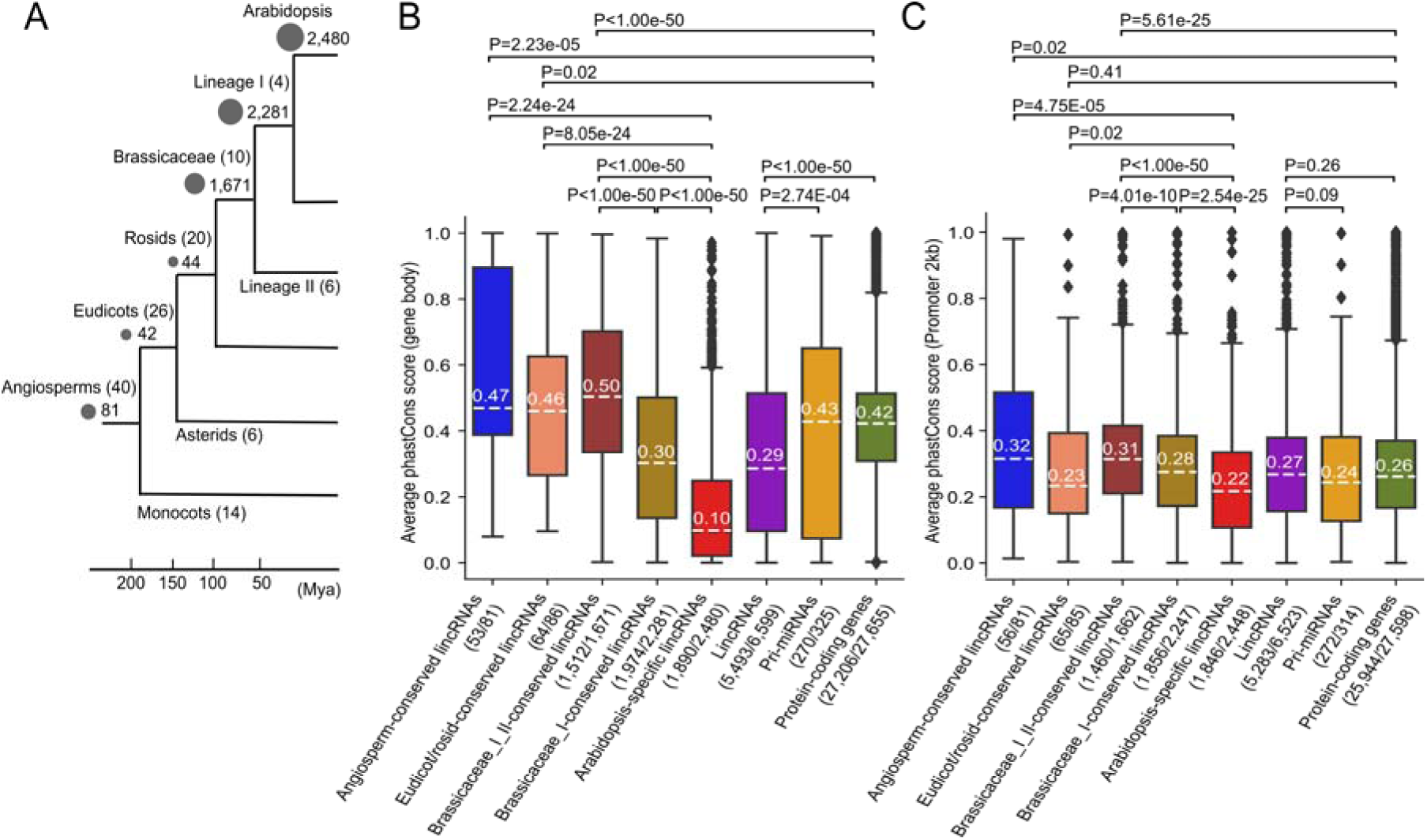
Sequence conservation analysis for lincRNAs of different evolutionary age categories. **(A)** Simplified species tree reporting the number of lincRNAs found for the different age categories, which are also indicated by the grey circle sizes. Numbers in parenthesis report the number of genomes included per clade to assess sequence similarity and define a lincRNA’s age category. Boxplot showing the average phastCons score for **(B)** gene bodies and **(C)** promoter regions (2kb upstream of transcription start site) of different lincRNA age categories and gene types (lincRNAs, pri-miRNAs, protein-coding genes). The numbers in parentheses report the number of gene bodies and promoters with at least 50% of informative nucleotides over the total number of gene bodies and promoters in that category, respectively. PhastCons score ranges from 0 (not under selection) to 1 (strong negative selection). P-values for pairwise Mann– Whitney *U* test are shown using the horizontal lines connecting the series.

The monotonous decrease in the number of lincRNAs for the older age categories suggests there is a continuous birth of lincRNA loci at the species level, with only a small fraction showing deep conservation in other plant families or orders.

Apart from detecting lincRNA homologs, we also evaluated evolutionary selection of lincRNA loci, pri-miRNAs, where globally the 21-24 miRNA sequence is the only conserved region at the nucleotide level, and protein-coding genes using phastCons scores (Siepel et al., 2005). This score reports the probability for each nucleotide to evolve neutrally or under negative, or purifying, selection (low or high phastCons score, respectively). In contrast to the age categories, which are based on finding similar homologous sequences in other plant genomes and do not give information about the mode of selection, phastCons works by fitting a two-state hidden Markov model to a genome-wide multiple sequence alignment and predicting, based on the pattern of nucleotide substitutions, conserved elements representing sites under purifying selection (Siepel et al., 2005). For lincRNA, pri-miRNA, and protein-coding gene loci, we compared phastCons scores for the gene body and the promoter region (2kb upstream of transcription start site) (Supplemental Data Set S2). In general, for gene bodies, lincRNAs are significantly less conserved than pri-miRNAs and protein-coding genes (Figure 2B, purple, yellow, and green series). However, we found that Angiosperm- and Eudicot/rosid-conserved lincRNAs were as conserved as protein-coding genes. Brassicaceae_I_II-conserved lincRNAs showed even significantly higher conservation levels than protein-coding genes, indicating high levels of purifying selection. Arabidopsis-specific lincRNAs show the lowest phastCons scores.

In contrast to the differences in gene body, the promoter scores are comparable for lincRNA, pri-miRNA, and protein-coding genes (Figure 2C, purple, yellow and green series). The level of purifying selection on promoter regions is similar in Angiosperm- and Eudicot/rosid-conserved lincRNAs as well as in protein-coding genes. We observed that conservation levels were again significantly higher in the Brassicaceae_I_II-conserved lincRNA promoters than protein-coding gene promoters. Taken together, these results indicate that Brassicaceae_I_II- conserved lincRNAs stand out in the level of purifying selection acting on these loci, both in the gene body and in their promoter, suggesting that the primary sequence of the RNA and its promoter regulation is a critical element for lincRNA function.

### Large-scale transcriptome analysis reveals highly-specific lincRNA expression in roots

Increasing evidence supports the tissue or cell-type specific role of lincRNA gene functions (Liu et al., 2012; Li et al., 2016; Cheng et al., 2017; Rai et al., 2019). To characterize spatiotemporal Arabidopsis gene expression patterns also considering lincRNAs, we curated, processed, and integrated 791 RNA-seq samples to construct a genome-wide gene expression atlas covering a wide range of tissues, developmental stages, and stress conditions (Supplemental Table S2 and Supplemental Data Set S3). To find a threshold of detectable expression above background, we normalized all data to establish detectable expression levels for protein-coding genes and lincRNAs (Ramskold et al., 2009; Li et al., 2016) (see Materials and Methods, Supplemental Figure S2A). A normalized transcripts per million (TPM) value >= 0.2 was considered to define 5,586 (84.6%) expressed lincRNAs and 284 (87.4%) pri-miRNAs, whereas a TPM >= 2 was used to define 26,254 (94.9%) expressed protein-coding genes. Globally, the expression breadth, defined as the number of samples in which a gene is expressed, was lower for lincRNAs (median: 12/791 samples) compared to pri-miRNAs (median: 35.5/791 samples) and protein-coding genes (median: 648/791 samples) (Figure 3A). Although the identification of lincRNAs is known to be impacted by the sequencing depth (Cabili et al., 2011; Liu et al., 2012; Sun et al., 2017), we did not observe a clear correlation between the expression breadth and sequencing depth (Pearson correlation coefficient, r=0.04, p=0.86) (Supplemental Figure S2B). This result indicates the reported expression values are not strongly biased by study-specific technical parameters. As previously reported, the expression level of lincRNAs is generally lower than that of pri-miRNAs and protein-coding genes (Figure 3B). The lincRNA loci part of the two “older” categories (Angiosperm and Eudicot/rosid-conserved lincRNAs) showed significantly higher expression levels compared with the three “younger” categories of lincRNAs (Brassicaceae_I_II-conserved, Brassicaceae_I-conserved and Arabidopsis-specific lincRNA). We also observed a six-fold difference in the fraction of expressed genes depending on their annotation source: lincRNAs annotated in only one resource where more frequently found as undetectable compared to lincRNAs annotated in two or more resources (20% versus 3%, respectively).

**Figure 3.**
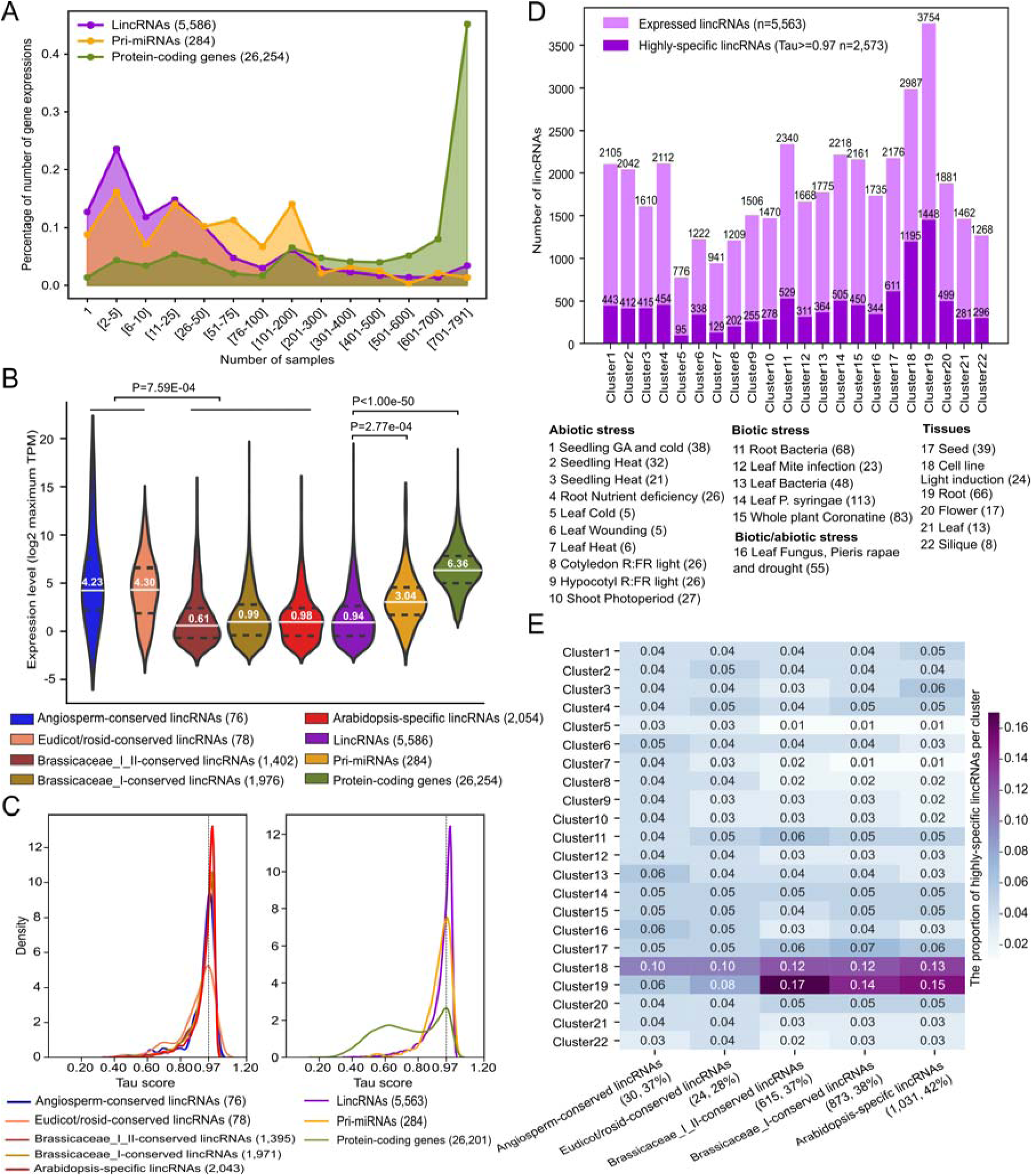
Expression analysis of lincRNAs. **(A)** Line chart showing the distribution of expression breadth for lincRNAs, pri-miRNAs and protein-coding genes across all samples **(B)** Distribution of the maximum TPM expression levels for different lincRNA age categories and gene types (lincRNAs, pri-miRNAs and protein-coding genes). **(C)** Distribution of tissue specificity tau scores for different lincRNA age categories and gene types. Tau scores range from zero to one, where zero means widely expressed, and one means very specifically expressed (detectable in only one cluster). The black dotted line represents a tau score of 0.97. **(D)** Bar chart reporting the number of expressed and highly-specific expressed lincRNAs for the different expression clusters. Cluster numbers and descriptions are shown below the chart, with numbers in parenthesis indicating the number of samples present per cluster. **(E)** Heatmap showing the proportion of highly-specific expressed lincRNAs in each cluster for each lincRNA age category. Cluster descriptions are the same as in panel D. Numbers in parenthesis report the number of genes per age category together with the fraction of lincRNAs showing highly-specific expression.

To identify groups of samples with similar expression patterns, 769 RNA-Seq samples were split into 22 expression clusters using meta-data curation (22 samples were discarded, see Materials and Methods). Globally, these clusters group Arabidopsis (Col-0) samples according to the organ or tissue considered and the stress, either biotic or abiotic, applied (Supplemental Data Set S3). As lincRNAs are proposed to exert their role in a tissue-specific manner (Liu et al., 2015), the tissue-specificity index (tau) (Yanai et al., 2005) for each lincRNA, pri-miRNA and protein-coding gene was calculated to estimate expression specificity between clusters (Figure 3C). LincRNAs were more specifically expressed than pri-miRNAs and protein-coding genes, with the median tau scores of 0.97, 0.95 and 0.74, respectively (P<0.001, Mann–Whitney U-test). Forty-six percent of all expressed lincRNAs (2,573/5,563) with a tau score ≥ 0.97 were defined as lincRNAs showing highly-specific expression (Supplemental Data Set S4). Even though we did not observe a uniform distribution of these highly-specific lincRNA over the 22 clusters (Figure 3D), especially cluster 19 (root), 18 (cell line light induction) and 3 (seedling heat) have the largest number of highly-specific lincRNAs. Between 28 and 42% of the highly-specific expressed lincRNAs were present in the different evolutionary categories, with the highest fraction in the Arabidopsis-specific lincRNAs (Figure 3E). Noteworthy, expression cluster 19, containing 66 samples covering both whole root and specific root cell types (Li et al., 2016), contains the highest fraction (14-17%) of highly-specific lincRNAs in the younger age categories (Figure 3E), hinting towards a role of these lincRNAs in root tissues.

To validate these tissue-specific expression patterns, we verified the expression of previously characterized Arabidopsis lincRNAs (Supplemental Table S3). The lincRNA *SVALKA*, which was identified in a cold-sensitive region of the Arabidopsis genome, was maximally expressed in the cold stress-related cluster 5 in our study (Kindgren et al., 2018). In addition, the lincRNA *MARS*, which is involved in the response to abscisic acid, seed germination and root growth under osmotic stress (Roule et al., 2022b), was found to be expressed at high levels in root cluster 19 as well as cluster 11 (root bacteria) and cluster 17 (seed). The lincRNA *ASCO*, reported to be involved in lateral root formation and response to pathogens (Bardou et al., 2014; Rigo et al., 2020), was found to be widely expressed, including cluster 11, which contained samples reporting bacterial flagellin stress responses. The Arabidopsis lncCOBRA1, also conserved in *Brassica rapa* (Kramer et al., 2022) and playing a role in seed germination, was found in our set of Brassicaceae_I_II-conserved lincRNA and showed the highest expression in cluster 17 (seed) and cluster 3 (seedling heat).

Among the highly-specific lincRNAs, the vast majority (66%) were derived from one resource, with lincRNAs identified in Liu et al. 2012 being most abundant (30%, Supplemental Figure S3A-B). Taken together, these findings indicate that the expression clusters offer a good starting point for the context-specific characterization of known and novel lincRNAs. Furthermore, young age categories, including the Brassicaceae-conserved and Arabidopsis-specific lincRNAs, show a bias for expression specificity in the root samples, with more than half (1,448/2,573) of the highly-specific lincRNAs active in root, suggesting a potentially diverse role of lincRNA networks in root growth and development.

### Experimental TF-lincRNA regulatory networks reveal active and complex gene regulation of Arabidopsis lincRNA genes

To integrate lincRNAs into epigenetic and transcriptional networks, we compared the chromatin states inferred by (Liu et al., 2018; Hazarika et al., 2022) for the different lincRNA gene sets delineated in our study. These chromatin states report a combination of multiple epigenetic marks along the genome and offer detailed insights in the locations and functions of regulatory regions and genes. Globally, lincRNAs show enrichment for vastly different chromatin states compared to protein-coding genes (Figure 4A). Chromatin states 33-34, typically associated with DNA methylation, repressive histone modifications and transposable elements, were strongly overrepresented in angiosperm-conserved lincRNAs, and to a lesser extent in lincRNAs showing highly-specific expression patterns. Chromatin states 31-32, also associated with DNA methylation and repressive histone modifications were strongly overrepresented in Arabidopsis-specific lincRNAs. Interestingly, these repressive chromatin marks were less enriched in the Brassicaceae-I_II-conserved lincRNAs, where chromatin states 13, 19 and 20 were most strongly overrepresented. State 13 refers to Polycomb group mediated deposition of trimethylation of the lysine 27 of histone H3 (H3K27me3) while states 19-20 denote accessible chromatin and TF ChIP binding. Chromatin states 19-20, but also states 31-32 and 34, were enriched in highly-specific lincRNAs. Although we found no evidence of a correlation between lincRNA age categories and chromatin states, our results revealed that a significant number of Brassicaceae_I_II-conserved lincRNAs have epigenetic signatures associated with Polycomb regulation and TF binding in accessible chromatin.

**Figure 4.**
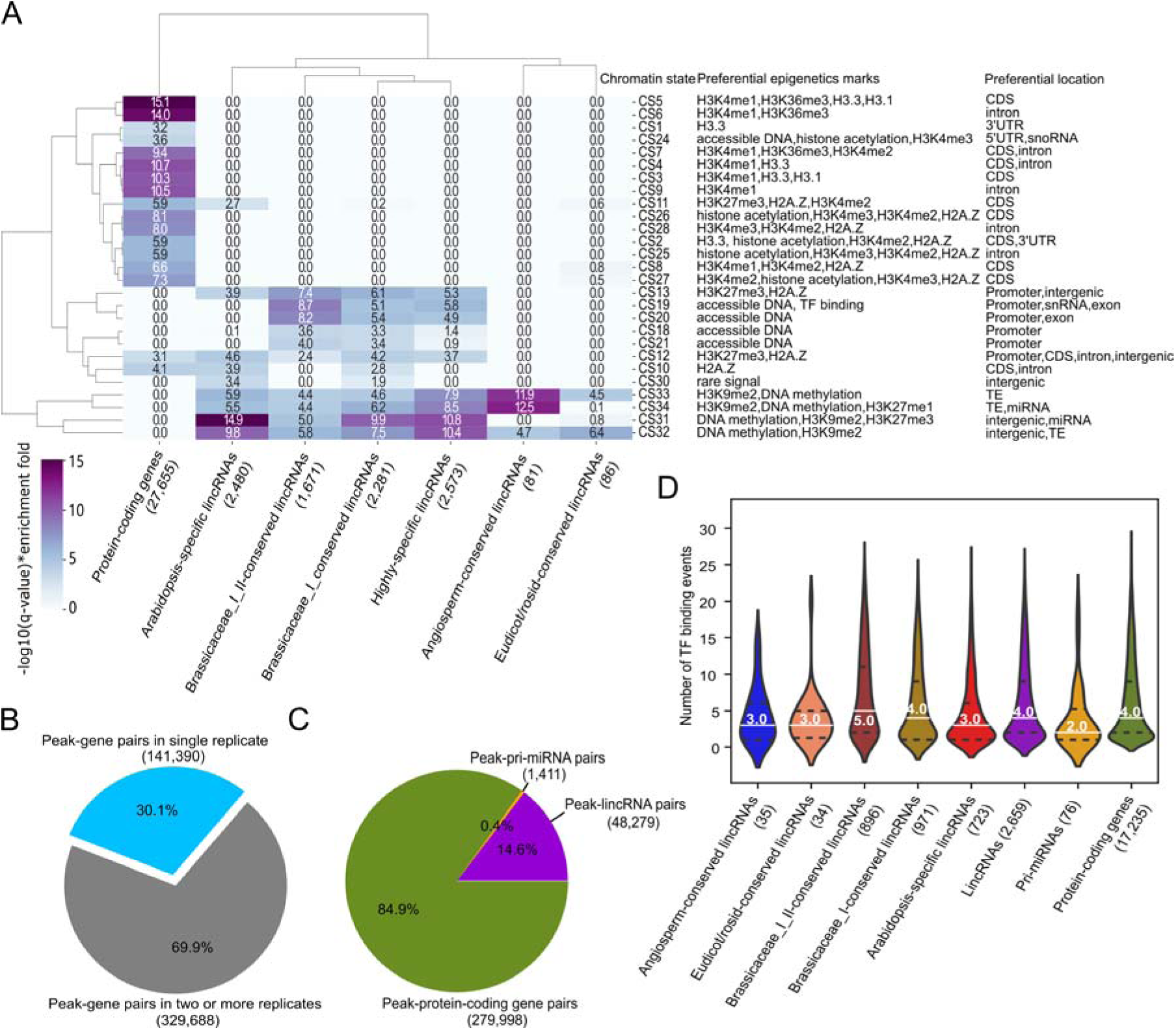
Chromatin state and TF ChIP-Seq peak annotation for different gene types. **(A)** Dendrogram showing the enrichment for different lincRNA gene sets (x-axis) towards different chromatin states (CS) (y-axis). The values report the product of -log10(q-value) and the enrichment fold. Only significant enrichment values are reported (q-value < 0.05). **(B)** The proportion of peak-gene pairs present in single replicate (blue) and two or more ChIP-Seq replicates (grey). **(C)** The percentage of three gene types assigned to peaks in two or more ChIP-Seq replicates. **(D)** Distributions of the number of TF binding events for lincRNA evolutionary age categories and gene types (lincRNAs, pri-miRNAs and protein-coding genes).

Based on the specific expression profiles for different lincRNA genes, the chromatin state information, as well as the high levels of promoter conservation, we next integrated TF chromatin immunoprecipitation (ChIP) data to further identify the regulators controlling lincRNAs. Before we used TF ChIP data to characterize the organization, complexity, and evolution of TF binding for protein-coding genes (Heyndrickx et al., 2014), hence we here re-processed publicly available ChIP-Seq to identify TF binding events potentially controlling lincRNA gene expression. A total of 114 TF ChIP-Seq datasets, covering 45 TFs with at least two biological replicates, were reprocessed (Supplemental Table S4). Starting from our genome annotation containing protein-coding genes, pri-miRNAs and the high-confidence lincRNAs, ChIP-Seq peaks were assigned to the closest genes. A gene was defined as a potential target gene for a profiled TF if it was the closest to at least one ChIP-Seq peak of that TF (see Material and Methods). Based on all 471,078 TF peak-target gene pairs identified from the 114 ChIP-Seq datasets, 329,688 (70%) of the peak-gene pairs were confirmed by two or more replicates and were kept to construct a robust TF- target gene regulatory network (Figure 4B). While most TF peaks were associated with protein-coding genes, we identified 48,279 (14.6%) peaks that were associated with lincRNAs (Figure 4C). Given the strong localization bias for TF binding in Arabidopsis (Heyndrickx et al., 2014; Yu et al., 2016), we only considered expressed target genes localized relative to the peak midpoint within a 2kb window (Supplemental Figure S4A), retaining 93.0% of the binding events (15,686 TF-lincRNA interactions, see Supplemental Data Set S5). Globally, 2,659 expressed lincRNAs had one or more TF binding events. Twenty-two out of the 45 TFs, belonging to the bHLH, HD-Zip, bZIP, C2H2, GRAS, MYB, NAC, NF-YB and NF-YC TF families, were each associated with at least 200 lincRNAs (Supplemental Table S5).

(Haudry et al., 2013) identified over 90,000 conserved non-coding sequences (CNS) in Arabidopsis that show evidence of transcriptional and post-transcriptional regulation. Comparing these CNS with the ChIP-Seq peaks of target genes revealed that most of the TF binding events close to lincRNAs (78.1%) and protein-coding genes (73.7%) overlapped with a CNS (Supplemental Figure S4B). This fraction was much lower for pri-miRNAs (57.0%). The highest fraction of ChIP-Seq peaks containing a CNS was detected for Brassicaceae_I_II-conserved lincRNAs loci (91.8%), indicating that these TF binding events are evolutionary constrained and potentially functional. Considering the different gene types, the median number of TF binding events per locus was higher for protein-coding genes and lincRNAs (median of 4 TFs) compared to pri-miRNAs, suggesting these genes are differently controlled (Figure 4D). Brassicaceae-conserved lincRNAs have more TF binding events than lincRNAs in any other age categories (Figure 4D). Furthermore, we observed a positive correlation between TF binding frequency and the conservation of lincRNA gene body (r=0.27, P=8.11e-92) (Supplemental Figure S4C). More precisely, a large faction of lincRNAs without any TF binding event also has very low phastCons scores, close to zero, indicating that these genes are not under purifying selection. We observed a neglectable correlation between the number of TF binding events and tissue specificity (r=0.02, p=1.69e-01). Altogether, lincRNAs experiencing stronger levels of purifying selection are bound, and potentially regulated, by more TFs, independent of their tissue specificity (Supplemental Figure S4D). Taken together, we generated a multi- level genome-wide characterization covering chromatin state information, promoter conservation, ChIP-based TF binding and CNSs, for all detectable lincRNA across >700 expression samples, permitting to rapidly define the biological context and relevance of lincRNAs in Arabidopsis regulatory networks.

### MYB44, PIF4 and KAN1 regulate Arabidopsis lincRNAs in different root cell types

To identify TF-lincRNA regulatory interactions active in specific cellular contexts, we used the previously defined expression clusters to combine TFs regulation and lincRNA expression in specific organs, tissues, or stress conditions. Considering all expressed lincRNAs and all TF peak-based regulatory interactions described above, we tested if specific expression clusters are overrepresented for lincRNAs controlled by specific TFs (Y-axis and X-axis in Figure 5A-B, respectively). We observed a significant overrepresentation for KANADI1 (KAN1) binding to lincRNA loci in 11 clusters, of which clusters 19 (root), 18 (cell line light induction), 14 (leaf *P. syringae*) and 6 (leaf wounding) showed the most significant overlaps (Figure 5A). Comparing the different clusters revealed that root cluster 19 contained numerous enriched TFs, including Arabidopsis Zinc-Finger protein 1 (AZF1), JAGGED (JAG), Phytochrome-Interacting Factor 5 (PIF5), Repressor of GA (RGA), PIF1, MYB3, KAN1 and HAT22. The observed patterns of TF-binding in different expression clusters were unique for lincRNAs and highly dissimilar compared to TF binding enrichment for protein-coding genes (Supplemental Figure S5A).

**Figure 5.**
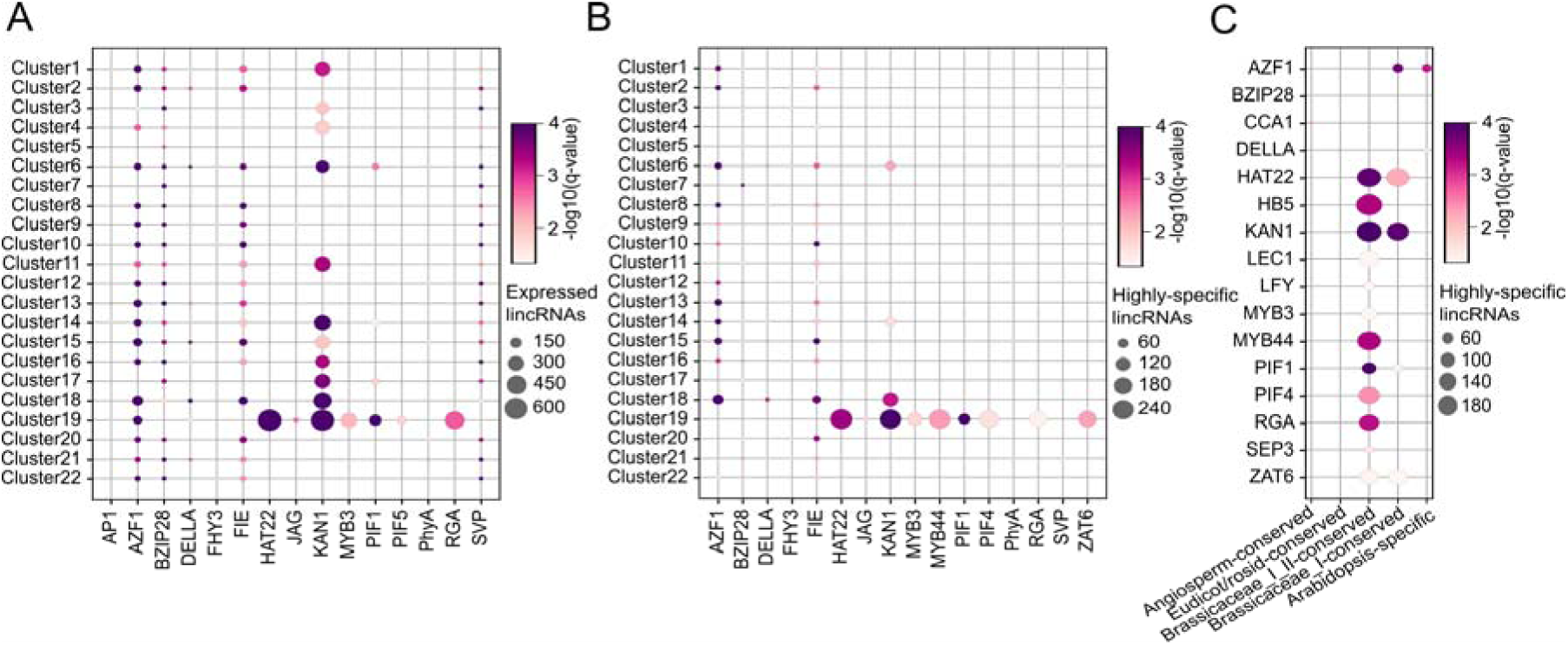
Overview of TF-lincRNA regulatory interactions in different expression clusters and age categories. The dot sizes represent the number of the lincRNAs while the color represents the statistical significance. Cluster descriptions are the same as in Figure 3D. **(A)** Bubble chart showing the enrichment of TF binding for expressed lincRNAs in different expression clusters. TFs lacking significant enrichment in any of the 22 clusters are not shown. **(B)** Bubble chart showing the enrichment of TF binding for highly-expressed lincRNAs in different expression clusters. TFs lacking significant enrichment in any of the 22 clusters are not shown. **(C)** Bubble chart showing the enrichment of TF binding for expressed lincRNAs in different age categories.

Focusing on the set of 2,573 lincRNAs showing highly-specific expression, most of the TF enrichments for root cluster 19 remained (Figure 5B). PIF1, PIF4, KAN1, HAT22, MYB3, MYB44, ZAT6 and RGA showed the largest overlap and all these TFs, apart from MYB3 and PIF1, contained >160 lincRNA target genes (Figure 5B and Supplemental Table S6). We confirmed that these eight TFs were all expressed in one or more samples of the root expression cluster 19 (Supplemental Figure S5B) and earlier studies have reported that these TFs, apart from HAT22, are involved in root development in Arabidopsis (Hawker and Bowman, 2004; Devaiah et al., 2007; Moubayidin et al., 2016; Tominaga-Wada and Wada, 2016; Zhao et al., 2016; Li et al., 2022). Comparing TF binding for the different age categories revealed that the Brassicaceae_I_II-conserved lincRNAs were most strongly enriched for these TFs (Figure 5C).

To validate the ChIP-based regulatory interactions for the identified TFs, we used reverse transcription quantitative PCR (RT-qPCR) analysis in roots of lines affected in TF expression such as overexpression or T-DNA inactivation (called TF perturbation lines). We used an inducible line for KAN1 (*KAN1-GR*), overexpression lines for MYB4 and PIF4 (*35S::MYB4*, 35S*::PIF4*) or quadruple mutant of the *pif1*, *pif2*, *pif3*, and *pif4* (*pifq*), and a knockout mutant for RGA (*rga28* T-DNA line). We then selected 27 potentially regulated lincRNAs showing high expression levels in the root cluster 19 and that were targeted by several of the selected TFs. For example, *LincRNA5331* and *LincRNA1119* were predicted to be regulated by the four TFs. We could detect deregulation for *LincRNA5331* and *LincRNA1119* in *35S::MYB44* and *pifq* roots, whereas the former was also deregulated in *KAN1-GR* and the latter in *35S::PIF4* (Figure 6A, Supplemental Figure S6). Overall, out of the 74 inferred regulatory interactions investigated, 36 were confirmed, meaning that for the tested TFs, a significant deregulation of the lincRNA was found in comparison to the control (“Has a peak and is DE”, in Table 1, Supplemental Table S7). For 23 out of the 27 tested lincRNAs, we confirmed one or more regulatory interactions (Figure 6A). The precision (i.e. the positive predictive value), varied between 27-65% while the average accuracy (i.e. the proportion of correct predictions among all genes examined) was 59%. For eight interactions the TF peak annotation to the lincRNA was unclear (“Has a putative peak and is DE” in Table 1), meaning these deregulated lincRNAs might also resemble confirmed interactions. Lastly, while the 10 interactions where we observed deregulation in the absence of a peak could indicate false predictions, they might also represent cases of indirect regulation controlled by the perturbed TF, influencing the expression of the profiled lincRNAs.

**Figure 6.**
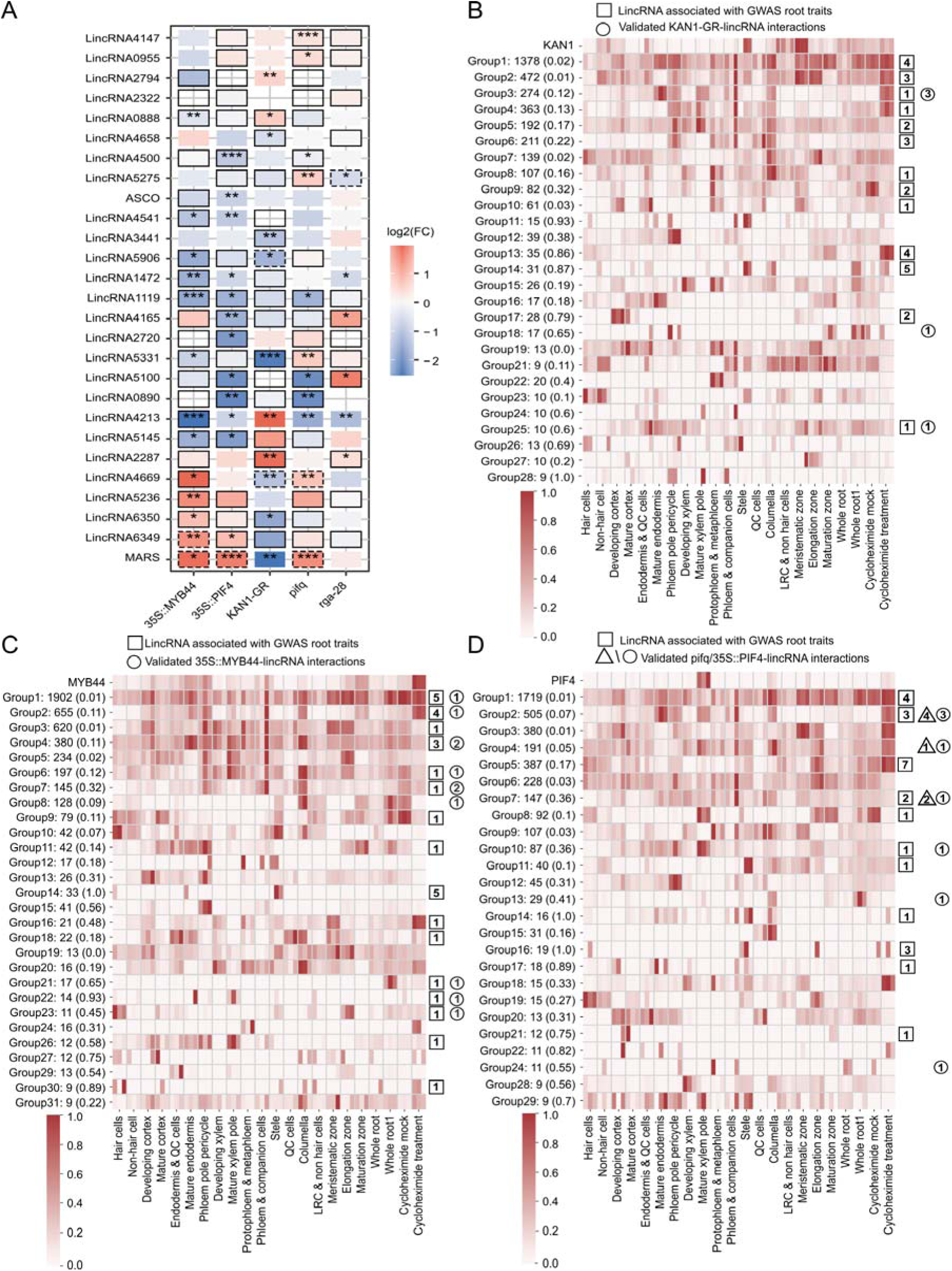
Experimental validation and characterization of TF-lincRNA regulatory interactions. **(A)** Heatmap showing log2 fold change (FC) of lincRNA relative expression levels in transcription factor (TF) overexpressing lines (35S::*MYB44*, 35S::*PIF4* and *KAN1-GR*) or TF knockout lines (*rga28* and *pifq* (*pif1/pif2/pif3/pif4* quadruple mutants)) vs. control wild-type lines in 14-day old roots. Expression values were determined by RT-qPCR. Asterisks indicate statistically significant differences (*p ≤ 0.05, **p ≤ 0.01, ***p ≤ 0.001) in an unpaired two-tailed Student’s t-Test (n =3). Solid boxes indicate TF ChIP peaks <2kb from the lincRNA gene. Dashed boxes indicate TF ChIP peaks were identified only in one ChIP-Seq replicate or the lincRNA is not the closest gene to the TF peak. **(B)** Heatmap showing KAN1 ChIP-Seq peak-based target genes in the cluster 19 root expression atlas. The first line shows the expression profile of the TF under investigation while the other lines show groups of target genes with similar expression. Next to the group number, the number of target genes is reported and in parenthesis the fraction of lincRNAs per group is indicated. The numbers in squares and circles report the number of GWAS root-related and RT-qPCR experimentally validated lincRNA genes, respectively. **(C)** Heatmap showing MYB44 ChIP-Seq peak-based target genes in the cluster 19 root expression atlas. Group information is the same as in panel B. **(D)** Heatmap showing PIF4 ChIP-Seq peak-based target genes in the cluster 19 root expression atlas. Group information is the same as in panel B, but here triangles and circles report the RT-qPCR experimentally validated lincRNA genes for the *pifq* and *35S::PIF4* lines, respectively.

**Table 1.**
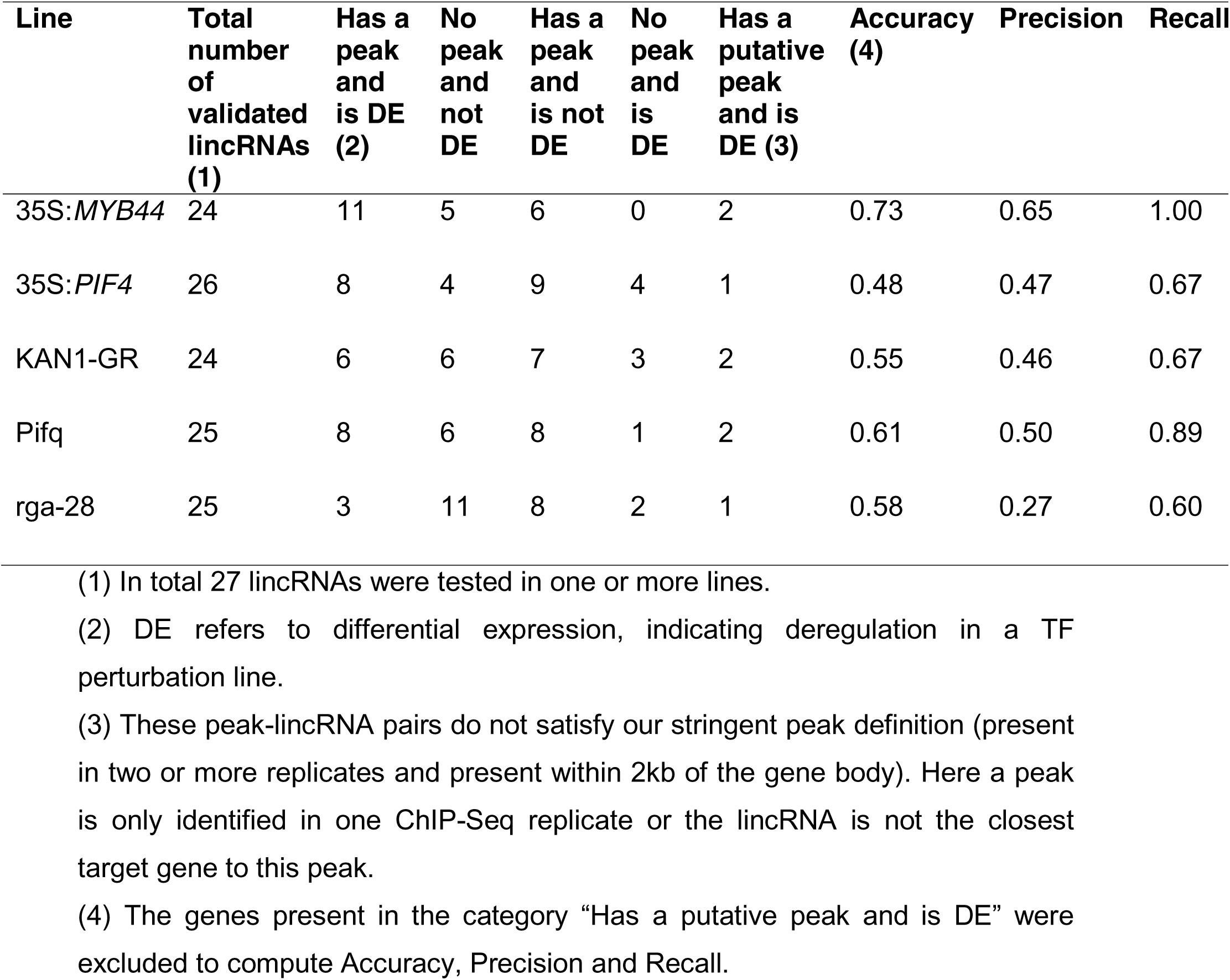
Summary of qPCR validation for TF-lincRNA gene pairs.

| Line | Total number of validated lincRNAs (1) | Has a peak and is DE (2) | No peak and not DE | Has a peak and is not DE | No peak and is DE | Has a putative peak and is DE (3) | Accuracy (4) | Precision | Recall |
| --- | --- | --- | --- | --- | --- | --- | --- | --- | --- |
| 35S:MYB44 | 24 | 11 | 5 | 6 | 0 | 2 | 0.73 | 0.65 | 1.00 |
| 35S:PIF4 | 26 | 8 | 4 | 9 | 4 | 1 | 0.48 | 0.47 | 0.67 |
| KAN1-GR | 24 | 6 | 6 | 7 | 3 | 2 | 0.55 | 0.46 | 0.67 |
| Pifq | 25 | 8 | 6 | 8 | 1 | 2 | 0.61 | 0.50 | 0.89 |
| rga-28 | 25 | 3 | 11 | 8 | 2 | 1 | 0.58 | 0.27 | 0.60 |

(1) In total 27 lincRNAs were tested in one or more lines.
(2) DE refers to differential expression, indicating deregulation in a TF perturbation line.
(3) These peak-lincRNA pairs do not satisfy our stringent peak definition (present in two or more replicates and present within 2kb of the gene body). Here a peak is only identified in one ChIP-Seq replicate or the lincRNA is not the closest target gene to this peak.
(4) The genes present in the category “Has a putative peak and is DE” were excluded to compute Accuracy, Precision and Recall.

Based on the experimental validation results confirming that several of the root-specific lincRNAs are controlled by the TFs inferred using the ChIP-based peaks, we next integrated genotype-phenotype relationships from the AraGWAS catalog. The aim of this analysis was to verify if root-related phenotypes have been reported for regions containing lincRNAs in genome-wide association studies (GWAS). After processing the significant associations from all GWAS studies present in the catalog and only retaining associations overlapping with lincRNA gene bodies or 2kb promoter regions (see Material and Methods), we identified 2,865 single nucleotide polymorphisms (SNPs) overlapping with 1,124 lincRNAs, covering 147 different studies. While 61.6% of the lincRNAs had only significant associations via SNPs in the promoter region, 38.4% had associations via SNPs in the gene body (194 and 238 lincRNAs with significant associations only in the gene body and in both gene body and promoter, respectively). After parsing and summarizing phenotype information from the available trait ontology annotations, we could link 23 lincRNAs to abiotic stress-related traits, 133 lincRNAs to flower-related traits, 33 lincRNAs to leaf-related traits, 376 lincRNAs to root-related traits, and 27 lincRNAs to seed-related traits (see Supplemental Data Set S6). The number of root-related GWAS annotations was much larger for lincRNAs targeted by TFs showing significant enrichment for binding in root cluster 19, compared to TFs lacking this enrichment (Supplemental Figure S7), confirming the functional relevance of these lincRNAs in roots. Overall, the reprocessed GWAS data indicate that hundreds of regions overlapping with lincRNAs loci are significantly associated with different plant traits and can be used to prioritize lincRNAs likely involved in specific biological processes.

To further investigate the relevance of the identified regulatory network, we focused on KAN1, MYB44, and PIF4, as these regulators had several root-expressed lincRNAs target genes that showed differential expression in the TF perturbation lines (Table 1). Starting from the protein-coding genes and lincRNAs expressed in root cluster 19 and annotated with a TF ChIP-seq peak, we first grouped target genes with similar expression in the root expression atlas. Secondly, for each group we determined the fraction of lincRNAs and verified if any of these were overlapping with our RT-qPCR experimentally validated lincRNAs or had root-related traits in the GWAS catalog. Finally, we verified if the tissue- or cell-type specific expression of the different groups was in agreement with the expression of the regulatory TF. For KAN1, we identified 14 groups that fulfilled these criteria (Figure 6B), of which eight contained two or more lincRNAs with root-related traits and several showed strong cell-type specific expression. Group 14, showing high expression in stele contained 27 lincRNAs (87%) of which five were associated with GWAS root traits. The high expression of KAN1 in the meristematic zone agreed with groups 1-2, 8-10, and 25, which together contained 12 confirmed lincRNAs, either through root-related traits or our experimental validations. For MYB44, we identified multiple groups containing GWAS root-related lincRNAs showing specific expression patterns in different cell types: group 4 contained multiple confirmed target genes expressed in mature endodermis while group 7 contained 3 confirmed lincRNAs (one GWAS root-related and two experimentally validated genes) showing high expression in phloem and companion cells (Figure 6C). For PIF4, which showed highest expression in mature xylem pole, groups 8 and 10 contained confirmed lincRNA target genes showing similar cell-type specific expression (Figure 6D). While other groups contained GWAS root-related lincRNAs showing strong expression in stele (groups 11, 14 and 16), our expression data does not confirm that PIF4 is actively expressed in the root stele. All GWAS root-related (58) or experimentally validated (21) lincRNAs regulated by KAN1, MYB44 or PIF4 can be found in Supplemental Table S8.

To confirm the root and cell-type specificity of the TF-lincRNA regulatory interactions, we integrated publicly available chromatin accessibility datasets of Arabidopsis roots based on Assay for Transposase-Accessible Chromatin using sequencing (ATAC-seq): three bulk datasets (Maher et al., 2018; Potter et al., 2018; Tannenbaum et al., 2018) and one single-cell dataset (Dorrity et al., 2021). Assessing the tissue specificity of our TF-lincRNA regulatory network revealed that 93% of the TF peaks associated with lincRNAs were detected in root accessible chromatin regions (ACRs), of which 21% were specifically detected in cell-type specific root ACRs. Randomly shuffling the TF peaks associated with lincRNAs showed that the observed overlap of TF-lincRNA regulatory interactions with bulk and single-cell root ACRs was 10-fold and 36-fold higher than expected by chance, respectively, confirming the high specificity of the inferred regulatory interactions. Of the 105 regulatory interactions covering GWAS root-associated or experimentally validated lincRNAs regulated by KAN1, MYB44, or PIF4, 97 were detected in the bulk root ACRs, of which 22 were detected in root cell-type specific ACRs. For several lincRNAs the cell-type specificity identified using the cell-type gene expression profiles were confirmed by the single-cell ATAC data (Supplemental Figure S8, Supplemental Table S8). For example, *LincRNA1736*, regulated by KAN1, and *LincRNA1296* and *LincRNA6434*, regulated by MYB44, were expressed in stele (xylem/phloem) and confirmed by xylem-specific ACRs. *LincRNA4180*, regulated by PIF4, was expressed in endodermis and cortex, which was confirmed by endodermis- and cortex-specific ACRs.

This analysis further supports that KAN1, MYB44 and PIF4 are controlling lincRNA genes showing highly specific expression in different root tissues and cell types. Additionally, the overlap with genetic associations hints to a role for many of these TF-controlled lincRNA loci in root growth and development.

## DISCUSSION

### Comprehensive annotation of Arabidopsis lincRNAs using transcriptomics and evolutionary genomics

In contrast to protein-coding genes, the characterization of lincRNAs is more challenging as we lack highly curated annotations and extensive experimental observations. Furthermore, the low levels of sequence conservation for the majority of lincRNA loci makes it difficult to translate biological knowledge learned in one species to another. Apart from co-expression network analysis, reporting putative (in) direct associations between lincRNAs and other genes, information about TF regulation of lincRNAs is scarce. Embedding lincRNAs in biological networks has great potential to define lincRNAs linked to specific cellular or morphological phenotypes.

Through the integration of eight lincRNA annotation resources, as well as mapping various conservation, chromatin, and expression features, we presented a global view on gene regulation for 5,586 expressed Arabidopsis lincRNAs. We strongly focused on using replicated samples when processing high-throughput datasets to obtain high-confidence gene information (Ponting and Haerty, 2022). Combined with comparative genome analysis yielding information about age categories and selection acting on gene bodies and promoter regions, we found that different subsets of lincRNAs have distinct molecular properties. While the high tissue-specificity and low levels of primary sequence conservation corroborate previous findings about lincRNAs (Necsulea et al., 2014; Ma et al., 2019; Palos et al., 2022), the analysis of lincRNA expression using a genome-wide gene expression atlas covering 769 samples revealed that lincRNA expression is widespread in different organs and not restricted to stress conditions, which is in agreement with previous studies (Jha et al., 2020; Corona-Gomez et al., 2022). In a recent study, Corona-Gomez and colleagues reconstructed a co-expression network to annotate lncRNAs with associated protein-coding genes. They identified several modules associated with root development or root-related stress functional annotation (Corona-Gomez et al., 2022). These results are consistent with the high representation of lincRNAs exhibiting highly-specific expression in the root expression cluster (unique set of 1448 lincRNAs), which is the highest number among all expression clusters we studied. When intersecting the different age categories with the expression clusters, Brassicaceae_I_II-conserved lincRNAs covered a large fraction of these root-expressed lincRNAs. While recent studies also identified a large number of context-specific lincRNAs expressed in root tip or meristem (Corona-Gomez et al., 2022; Palos et al., 2022), our age category analysis revealed that many lincRNAs that originated in the common ancestor of Brassicaceae lineages I and II, showed specific expression in various root cell types.

We observed a clear trend of increasing levels of lincRNA gene expression when going from young to old age categories. However, quantifying selection levels acting on different loci revealed that the oldest age categories, showing the most wide-spread and highest expression, are not undergoing the highest levels of purifying selection. While older categories like angiosperm-conserved and Eudicot/rosid-conserved lincRNAs had similar median phastCons scores as protein-coding genes, Brassicaceae_I_II-conserved lincRNAs showed the highest levels of purifying selection, both in their gene body and promoter. In animals, a significant increase of dynamically expressed genes and higher levels of purifying selection has been reported for older age categories (Necsulea et al., 2014; Sarropoulos et al., 2019). Furthermore, massively parallel reporter assays surveying thousands of human promoters revealed that tissue-specific lincRNAs had fewer TF motifs compared to ubiquitously expressed genes (Mattioli et al., 2019). In contrast to Arabidopsis protein-coding genes, this pattern was not found for lincRNAs in our analysis (correlation between tau score and TF binding frequency = 0.02). These results suggest, based on the genome-wide TF binding data available in Arabidopsis, that the complexity in TF control of lincRNAs is different between plants and animals and that the observed pattern of highly-specific expression and purifying selection for Brassicaceae_I_II-conserved lincRNAs deviates from the global trends observed in animals. Therefore, these unique properties suggest that these lincRNAs, which only emerged 42 million years ago, are better integrated in plant networks when compared to young lincRNAs in animals.

### Integrative regulatory annotation of Arabidopsis lincRNAs

The regulatory annotation using genome-wide chromatin state information revealed that protein-coding genes and lincRNAs have distinct chromatin signatures. The enriched chromatin states for lincRNAs were largely variable between, and sometimes within, the different age categories. While states associated with DNA methylation and repressive histone modifications were most strongly overrepresented in the youngest (Arabidopsis-specific) and oldest (angiosperm-conserved) lincRNA age categories, states denoting polycomb group mediated deposition of H3K27me3 and accessible chromatin were most enriched for Brassicaceae_I_II-conserved lincRNAs. H3K27me3 is a repressive covalent histone modification resulting from the activity of Polycomb repressive complexes. It was recently shown that a reduction in H3K27me3 levels leads to a decrease in the interactions within Polycomb-associated repressive domains, resulting in a global reconfiguration of chromatin architecture and transcriptional reprogramming during plant development (Huang et al., 2021). Chromatin accessibility is a hallmark of regulatory DNA as it allows sequence-specific binding of TFs, key components of transcriptional regulatory networks (Schmitz et al., 2022). The association of these chromatin states with specific sets of lincRNAs strongly indicates active transcriptional regulation.

To identify context-specific TF regulation potentially driving the highly-specific expression observed for many lincRNAs, we reprocessed, filtered and annotated 114 ChIP-Seq experiments covering 45 TFs, yielding a TF-lincRNA gene regulatory network containing 2,659 lincRNAs and 15,686 interactions. To assess the potential functionality of these inferred regulatory interactions, we overlapped CNSs identified using nine Brassicaceae genomes, which confirmed that TF peaks close to lincRNAs show similarly high levels of sequence constraint compared to protein-coding genes (74-78%). Furthermore, TF binding events close to Brassicaceae_I_II-conserved lincRNAs showed the highest levels of CNS conservation (92%), which agrees with the very high phastCons promoter scores we observed for this age category. While Palos and co-workers reported that CNSs significantly correlated with gene bodies of Brassicaceae-conserved lincRNAs (Palos et al., 2022), our results revealed that also lincRNA promoters and TF binding sites are strongly conserved for Brassicaceae_I-II-conserved lincRNAs. Ultraconserved CNSs, frequently associating with TF binding sites for key plant regulators controlling essential biological processes, have been identified for thousands of protein-coding genes (Van de Velde et al., 2016). Our results suggest that such deep conservation of cis-regulatory elements is extremely rare for lincRNAs, as only a small number of lincRNA genes show deep evolutionary conservation. Such deeply conserved CNSs for protein-coding genes frequently occur in divergent gene pairs, where they form mini-regulons representing conserved transcriptional units of co-regulated and co-expressed neighboring genes (Van de Velde et al., 2016). It is currently unclear if such conserved transcriptional regulons also exist for lincRNA loci and could explain the observed patterns of positionally-conserved but sequence-diverged lncRNAs (Mohammadin et al., 2015).

The integrated TF binding information showed that promotors of lincRNAs differ strongly from those of protein-coding genes, but also revealed high levels of heterogeneity among the different age categories. In animals, promoters of protein-coding genes contain more TF binding sites than those of lincRNAs, suggesting a stronger and more complex transcriptional regulation of the former (Necsulea et al., 2014; Sarropoulos et al., 2019). When comparing the number of TF binding events for the different gene types in Arabidopsis, we observed no difference in TF binding frequency for protein-coding genes and lincRNAs, indicating that lincRNAs are also regulated in a complex manner in plants. While no correlation between lincRNA expression tissue-specificity and TF binding frequency was found, a positive correlation between TF binding frequency and the level of purifying selection was observed. Again, Brassicaceae_I_II- conserved lincRNAs stood out having the highest number of binding TFs, suggesting that neither the age nor the expression of a lincRNA, but its importance for plant fitness, is a major factor in determining its regulatory complexity. While our results confirm that broadly expressed protein-coding genes, showing high expression breadth, are positively correlated with the number of regulating TFs (Heyndrickx et al., 2014), we did not observe this global trend for lincRNAs, indicating that the regulatory properties of TF control for plant protein-coding genes and lincRNAs are different. Although the number of Arabidopsis TFs profiled using ChIP-Seq may be considered as limited, future research will have to address whether the complexity of TF control varies for lincRNAs, as well as protein-coding genes, active in different organs, tissues, or stress conditions.

### A TF-lincRNA gene regulatory network identifies KAN1, MYB44 and PIF4 as regulators controlling root lincRNAs

Through integration of our spatiotemporal expression clusters and the TF- lincRNA gene regulatory network, we identified eight TF regulators showing a significant enrichment for TF binding close to lincRNAs specifically expressed in roots. While the overlap between TF ChIP-Seq peaks and CNSs gave an indirect indication of the potential functionality of TF binding sites close to hundreds of lincRNA loci, we experimentally validated a set of inferred regulatory interactions, focusing on 27 root-expressed lincRNAs and 5 TF perturbation lines. The number of confirmed regulatory interactions as well as the positive prediction values found for the tested TFs and lincRNAs here (27-65%) are 3 to 4 times higher than the fraction of TF-bound protein-coding genes also showing deregulation reported for a set of TF regulators involved in flowering (7-22%) (O’Maoileidigh et al., 2014). Compared to the discovery rates obtained for large-scale phenotypic screens of insertional lines (1.3%) (Ransbotyn et al., 2015), our discovery rates for deregulation are 20-50 fold higher. Globally, for 85% of the profiled lincRNAs one or more confirmed regulatory interactions were found. These findings indicate that the ChIP peak-based inference of TF regulation is a promising approach to characterize TF regulation of lincRNAs. Additional deregulated lincRNAs lacking a TF peak were also identified, which might be due to an indirect effect caused by crosstalk between different regulators in the Arabidopsis root as well as the type of mutations chosen in the perturbed TF lines (e.g. partners lacking in overexpressing lines, compensatory effects in gene families) (Heyndrickx et al., 2014).

While detailed functional characterizations of lincRNA genes are scarce, the integration of GWAS information allowed us to identify genetic associations for 1,124 lincRNAs covering various traits. While this number, corresponding to a frequency of 9%, is slightly lower compared to that of protein-coding genes associated with a specific trait in the AraGWAS catalog (3,030/27,655 = 11%), it does confirm the great potential of this largely untapped resource to biologically characterize lincRNAs potentially controlling different plant traits. As shown for KAN1, MYB44 and PIF4, multiple groups of co-expressed lincRNAs were identified bound by one of these TFs. Most of these groups showed strong cell- type specific expression and contained lincRNAs that were annotated with root- related traits. While most regulatory interactions were confirmed by TF peaks overlapping with root bulk ACRs, for several lincRNAs cell-type specificity was confirmed based on single-cell ATAC data, despite the high sparsity associated with this data type.

While co-expression networks and modules containing lincRNA genes cannot differentiate between direct and indirect regulatory interactions and lack functional information about individual lincRNAs (Corona-Gomez et al., 2022; Palos et al., 2022), our complementary approach relying on TF regulation and GWAS information overcomes some of these shortcomings. Taken together, the integration of different gene annotations combined with information about evolutionary conservation, selection, expression, TF regulation and GWAS data yielded new insights on the biological relevance of hundreds of Arabidopsis lincRNAs and offers a promising strategy to identify lincRNAs involved in different aspects of plant biology.

## MATERIALS AND METHODS

### Prediction of lincRNAs from in-house dataset

Paired-end RNA-seq datasets with high sequencing depth conducted in previous projects in the group at the Institute of Plant Sciences Paris-Saclay were used to predict additional lincRNAs. All data came from experiments carried out in the Col0 ecotypes and involved nsra/b mutant seedlings in response to NPA/NAA treatment GSE65717 and GSE116923 (Tran Vdu et al., 2016; Bazin et al., 2018), seedlings with modified expression of the *ASCO* lincRNA GSE135376 (Rigo et al., 2020), root tip submitted to a short phosphate starvation kinetic GSE128250 (Blein et al., 2020) and a lateral root initiation kinetic from a binding essay of five time points without replicates (6h, 12h, 24h, 36h and 48h after binding, unpublished). All reads were quality trimmed using Trimmomatic. For each library independently, cleaned reads were mapped on TAIR9 genome sequence with STAR (version 2.7.2a) using Araport11 as a guided annotation with the following additional parameters: --alignIntronMin 20 --alignIntronMax 3000. For each alignment file, StringTie (version 2.1.4) was used to predict transcripts using

Araport11 annotations as a guide (additional parameters: -c 2.5 -j 10). GFFcompare (v0.12.6) was then used to isolate the new transcripts in comparison to Araport11 gene annotation (removing transcripts with class code =, c, e or s). The different transcripts prediction were then combined using StringTie in merge mode (additional parameters: -F 0 -T 0 -c 0 -f 0 -g 0 -i). The final set of transcripts was compared against Araport11 annotation with GFFcompare removing all transcripts with a class code of =, c, e, s or m. Transcript were then associated with their already annotated gene or to newly defined genes in case they were predicted in unannotated portion of the genome. Coding potential was then assets using COME (Hu et al., 2017), Coding Potential Calculator CPC (v0.9-r2), CPC2 (Kang et al., 2017) and infernal (v1.1.2) (Nawrocki and Eddy, 2013) against Rfam v14.1 (Kalvari et al., 2021) with default parameters. Non-coding transcripts were the ones predicted by CPC, CPC2 and COME as non-coding and having no hits against tRNA, rRNA, snRNA or snoRNA genes in Rfam.

### Integration of lincRNA gene annotations

*Arabidopsis* lincRNAs were collected from public databases including Araport11 (Cheng et al., 2017), CANTATAdb (Szczesniak et al., 2016), NONCODEv5 (Fang et al., 2018), PLNlncRbase (Xuan et al., 2015), obtained from publications (Liu et al., 2012; Nelson et al., 2017; Zhao et al., 2018) or shared by Andrew D. L. Nelson from the Boyce Thompson Institute at Cornell using their previously published method, and the in-house dataset. These collections contain lincRNA annotations based on transcriptomic information coming from a wide variety of organs including seedling, root, pollen, rosette leaf, endosperm, seed, siliques, inflorescence, flower, floral buds, as well as abiotic stress treatments. The pipeline used for the identification of putative lincRNA is described in Supplemental Figure S1A. (1) LincRNA transcripts with a length of at least 200 bp were retained. (2) Only transcripts that were at least 500 bp away from any protein-coding gene were retained and considered as intergenic (Liu et al., 2012; Yamada, 2017). (3) Transcripts lacking strand information were discarded. (4)

The coding potential of transcripts was assessed using the Coding-Non-Coding Index (CNCI; Version 2) (Sun et al., 2013), CPC2, and Pfam-scan (PFAM) (Finn et al., 2016), and only those transcripts fulfilling the CPC2 (cutoff < 0), CNCI (cutoff < 0) and PFAM (E-value 1e-5) criteria were retained. (6) All candidate intergenic transcripts were assigned to lincRNA loci using the GFFcompare program (Pertea and Pertea, 2020). The number of transcripts retained after each filtering step is reported in Supplemental Figure S1B.

### Evolutionary conservation analysis

Our set of Arabidopsis lincRNAs was classified into distinct evolutionary age categories based on sequence similarity. The sequences of 6599 lincRNA loci were extracted using BEDTools getfasta v2.30.0 (Quinlan and Hall, 2010). Genome sequence data for 40 plant species were obtained from PLAZA 5.0 Dicots and PLAZA 5.0 Monocot (Van Bel et al., 2021), representing 26 eudicots (20 rosids and 6 asterids ) and 14 monocots. The 20 rosids contain 10 Brassicaceae species. LincRNA homologs were identified using sequence similarity searches against these 40 genomes using BLASTn (Altschul et al., 1990) and applying an E-value cutoff of 1e–10 (Nelson et al., 2017). Classification rules were defined to construct five evolutionary age categories. A lincRNA was deemed conserved in the Angiosperm evolutionary age category when at least one homolog was found in eudicots and one homolog in monocots. LincRNA was defined as a eudicot/rosid-conserved lincRNA with at least one homolog in rosids, one homolog in asterids, and no homolog in monocots, in addition with one homolog only in rosids. The lincRNA was assigned as a Brassicaceae_I_II-conserved lincRNA with at least one homolog in Brassicaceae lineage I, one homolog in Brassicaceae lineage II, and no homologs outside the Brassicaceae species. A lincRNA that had at least one homolog in a Brassicaceae lineage I species, apart from Arabidopsis, was defined as Brassicaceae_I-conserved lincRNA. The last category, defined as Arabidopsis-specific lincRNA, were restricted to Arabidopsis lincRNAs without homologs in any of the other species.

We downloaded the GFF annotation file for 27,655 protein-coding genes and 325 pri-miRNAs from the Araport11 genome release (Cheng et al., 2017) and extracted gene body sequences using BEDTools getfasta v2.30.0. We also extracted the promoter region of 2kb upstream of the transcription start site for lincRNAs, pri-miRNAs, and protein-coding genes. Noted that the promoter of a gene was shortened when it overlapped with a nearby gene sequence.

The sequence conservation of gene body and promoter region per gene type was evaluated using phastCons scores, which were calculated using the alignments of 20 angiosperm plant genomes (Hupalo and Kern, 2013). The phasCons scores were downloaded from the araTha9 genome browser available at genome.genetics.rutgers.edu as a bedgraph file. The bedgraph, consisting of variable width bin of equal phastCons score, was reprocessed to use a fix bin width of 1nt. In case of absence of a phasCons score on a portion of the genome, no bin was created which allow making the difference between nucleotides with a score (informative nucleotides) or absence of score (non-informative nucleotides). The average phastCons score of each annotation or promoter was computed using the BEDTools map v2.30.0 with the “–c4 –o mean” options, giving the average phastCons score using only informative nucleotides. The number of informative nucleotides of the genome proportion with phasCons score was computed using BEDTools intersect v2.30.0 Only for loci with at least 50% of informative nucleotides average phastCons scores were computed and reported in Figure 2 B-C (other loci were discarded).

### Expression analysis of lincRNAs

To generate an expression atlas for lincRNAs, pri-miRNAs and protein-coding genes, we used Curse (Vaneechoutte and Vandepoele, 2019) to search and curate relevant RNA-seq experiments. Details of the 18 RNA-seq experiments across all 791 samples are shown in Supplemental Table S2 and Supplemental Data Set S3. Next, we imported the expression metadata to the Prose tool (Vaneechoutte and Vandepoele, 2019), which downloads the raw data from SRA by using the SRA toolkit, performs quality control and adapter detection, trimmomatic to perform adapter clipping and quality trimming and finally Kallisto (Pertea et al., 2015) for quantifying transcript expression to normalized transcripts per million (TPM) values. We downloaded the transcript FASTA file for 27,655 protein-coding genes and 325 pri-miRNAs from the Araport11 genome release (Cheng et al., 2017) and retrieved transcript sequences for 6,599 lincRNAs using gffread (Pertea and Pertea, 2020). Gene-level atlases were created by summing the TPM values of all transcripts. We retained TPM value per gene across the biological replicates and took an average of TPM values per gene from technical replicates, resulting in an expression atlas covering 791 samples.

We used a simulation experiment (Ramskold et al., 2009; Li et al., 2016) to determine thresholds for detectable expression in TPM for protein-coding genes and lincRNAs. The protein-coding genes were used as true positives and the lincRNAs were used as true negatives. We calculated the false-positive rate and false-negative rate at different TPM thresholds for each sample. The applied cutoff of 0.2 TPM to define lincRNA and Pri-miRNAs expression is a good trade-off to detect lowly expressed lincRNAs and keep false-positives under control (false positive rate<0.10). A more stringent threshold TPM >= 2 was used to define the expression of protein-coding genes.

To identify the clusters containing samples with related gene expression information, a log_2_-transformed gene expression matrix (TPM+1) was used for t-SNE clustering with perplexity 30 and n_iter = 1000 (Pedregosa et al., 2011). 22 clusters were retained, the largest of which contained 113 samples and the smallest contained 5 samples. Twenty-two samples were not clustered and removed (Supplemental Data Set S3). Expression specificity per gene type, for the gene expression matrix described above, was measured using tissue-specificity index (tau), defined by the following equation (Yanai et al., 2005):

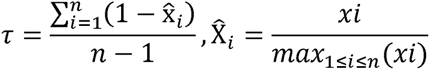

re *x_i_* was defined as the average TPM value of per cluster and *n* corresponds to the number of clusters analyzed.

Groups of genes showing similar expression profiles reported in Figure 6 B-D were identified using CAST clustering and only clusters with at least 10 genes were retained (Vandepoele et al., 2009).

### Chromatin states analysis and TF peak annotation

The enrichment of the lincRNA sets in different chromatin states (CSs) of Arabidopsis from (Liu et al., 2018) was done by shuffling the lincRNA genome coordinates 1000 times over the whole Arabidopsis genome. Then, we compared the distribution of the number of lincRNAs expected to overlap by chance with each CS with the real number of overlaps, and we used these values to calculate enrichment statistics: p-value as the number of times the real overlap was higher than the overlap with any of the 1000 shuffled lincRNA sets, and enrichment fold as the real overlap divided by the median of overlap expected by chance (median of the 1000 shuffling events). The p-value was adjusted for multiple testing using Benjamini-Hochberg correction (significance level 0.05). For visualization purposes, the two enrichment metrics were combined into the π-value, which is the -log10(p-value)*enrichment fold.

TF ChIP-Seq peak coordinates were retrieved from the PlantPAN 3.0 database (Chow et al., 2019) and (Song et al., 2016). The original ChIP-Seq of KAN1, MYB44 and PIF4 were derived from these studies (Merelo et al., 2013; Pfeiffer et al., 2014; Song et al., 2016). For each peak the closest gene was identified and only peaks confirmed in two or more replicates and within a 2kb window of the gene body were retained. Starting from all TF – target gene pairs (either a protein-coding gene or a lincRNAs; Supplemental Data Set S5), enrichment analysis was performed to identify enriched TFs in different expression clusters or age categories. For all enrichment analyses the hypergeometric distribution was applied and the q-value of enrichment was determined using the Benjamini– Hochberg correction for multiple hypotheses testing.

### Identification of GWAS-associated genes

GWAS data were collected from AraGWAS (Togninalli et al., 2020) and overlapped with the gene body and promoter 2kb sequences of lincRNAs to associate with the phenotype of interest. All significant associations (Permutation threshold <0.05 and FDR <0.05) were retained for screening for minor allele frequency>0.01, resulting in 1,124 lincRNAs, covering 147 different studies. These significantly associated lincRNAs were classified into five main traits by ontological annotation, including root, seed, flower, leaf, and abiotic-related traits (see Supplemental Data Set S6).

### Overlap with accessible chromatin regions

ATAC-seq data for Arabidopsis root hair, non-hair and whole roots were collected from three publications (Maher et al., 2018; Potter et al., 2018; Tannenbaum et al., 2018), and scATAC-seq data for Arabidopsis root epidermis, endodermis, stele (pericyle, xylem, phloem), and cortex cells were collected from (Dorrity et al., 2021). Cell type-specific marker peaks were identified by scATAC-seq data (p<0.05 and avg_lofFC>0). BEDtools intersect v2.30.0 (Quinlan and Hall, 2010) was used to detect whether TF ChIP-seq peaks associated with lincRNAs overlapped with ACR peaks. Only overlaps between ACRs and TF ChIP peaks in at least two replicates were retained, requiring that at least of 10% of the ChIP peak was covered by the ACR. BEDtools shuffle v2.30.0 (with parameter -chrom) was used to shuffle the TF ChIP-seq peaks associated with lincRNAs.

### Validation of TF-lincRNA regulatory interactions using reverse transcription-quantitative PCR

Total RNA from five to eight 14-day roots grown vertically on solid MS ½ media was extracted using TRI Reagent (Sigma-Aldrich) and digested with RNase-free DNase (Fermentas) following the manufacturer’s recommendations. cDNA was synthetized using Maxima Reverse Transcriptase (Thermo Scientific). Expression analysis by RT-qPCR was performed using SYBR Green master I (Roche) and the LightCycler® 96 system following a standard protocol (40 cycles, 60°C annealing). Data were analyzed using the ΔΔCt method with PP2A (PROTEIN PHOSPHATASE 2A SUBUNIT A3 (AT1G13320)) as reference transcript for normalization of RT-qPCR data. WT Col-0 plants grown at the same time were used as sample reference. For the dexamethasone inducible KAN1-GR expression lines analysis, after 14 days on MS media, the plants were transferred for one day either on dexamethasone containing plates (10µM) or only DMSO (dexamethasone solvent) as control. Primers used are listed in Supplemental Table S9. Three biological replicates were performed per condition. Statistical analyses were performed using the unpaired two-tailed Student’s t-Test (GraphPad prism).

## SUPPLEMENTAL DATA

**Supplemental Figure S1.** Integration of lincRNA resources.

**Supplemental Figure S2.** Definition of expressed genes and influence of sequencing depth.

**Supplemental Figure S3.** Resource annotation for highly-specific expressed lincRNAs.

**Supplemental Figure S4.** Regulatory properties of lincRNAs.

**Supplemental Figure S5.** TF binding and expression.

**Supplemental Figure S6.** RT-qPCR validation results.

**Supplemental Figure S7.** GWAS traits associated with lincRNAs confirmed by ChIP-Seq TF binding.

**Supplemental Figure S8.** Identification of cell type-specific TF-lincRNA regulatory interactions.

**Supplemental Table S1.** The number of homologous lincRNAs aligned to the genomes of 40 species.

**Supplemental Table S2.** Summary of RNA sequencing data.

**Supplemental Table S3.** List of lincRNAs with known functions.

**Supplemental Table S4.** Overview of TFs ChIP-Seq data used in this study.

**Supplemental Table S5.** TF and the absolute number of target genes per gene type.

**Supplemental Table S6.** Overview of TFs showing significant enrichment for binding to highly-specific expressed lincRNAs for the different expression clusters (corresponds to Figure 5B).

**Supplemental Table S7.** Overview of lincRNA peak annotation and RT-qPCR validation.

**Supplemental Table S8.** RT-qPCR confirmed/associated with GWAS root traits lincRNAs expression information for the different samples of expression cluster 19 (root) (corresponds to Figure 6B-D) and regulatory interactions detected in ACRs (corresponds to Supplemental Figure S8).

**Supplemental Table S9.** RT-qPCR primer sequences used in this study.

**Supplemental Data Set S1.** Full list of putative lincRNAs integrated from different resources.

**Supplemental Data Set S2.** PhastCons Scores for lincRNA loci and evolutionary age category.

**Supplemental Data Set S3.** Metadata for 791 samples of RNA-seq and cluster information.

**Supplemental Data Set S4.** The list of the highly-specific lincRNAs.

**Supplemental Data Set S5.** The table of TF-lincRNA interactions.

**Supplemental Data Set S6.** The lincRNA loci containing significant GWAS hits.

## AUTHOR CONTRIBUTIONS

L.L., M.C., T.B., and K.V., designed the research. L.L., T.D., and T.B., performed data analysis. N.M.P. performed chromatin state analysis. M.H. performed experimental work. L.L., T.B., and K.V., wrote the manuscript.

## ACKNOWLEDGEMENTS

We thank Mariek Dubois, Andrés Ritter, Tereza Vavrdova, Freya De Winter, Rebecca De Clercq for kindly providing the Arabidopsis lines. We thank Olivier Martin and Federico Ariel for providing comments on the manuscript. The IPS2 is benefited from the support of Saclay Plant Sciences-SPS (ANR-17-EUR-0007).

L.L. is supported by the China Scholarship Council for a PhD fellowship (201808530499). M.H. is supported by a PhD fellowship from the Fondation pour la Recherche Médicale (ECO202106013730).

