## Supplemental Figures for "Regulatory annotation identifies KAN1, MYB44 and PIF4 as regulators of Arabidopsis lincRNAs expressed in root"

A

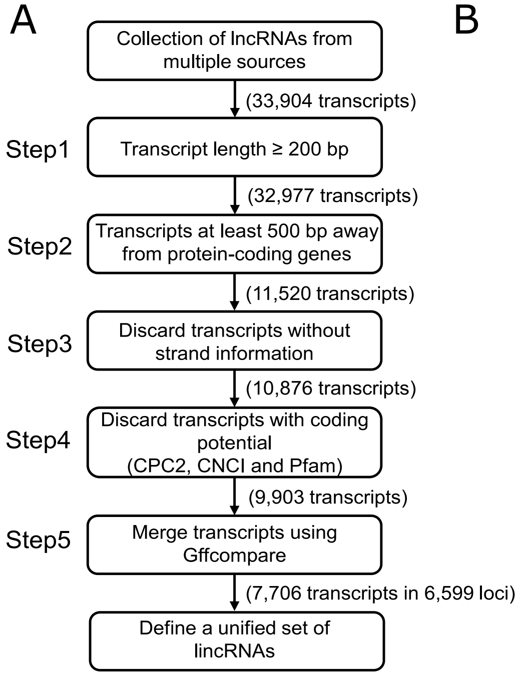


B

| Resource | No. raw transcripts | Step1 | Step2 | Step3 | Step4 | Reference |
|---|---|---|---|---|---|---|
| Araport11 | 2,455 | 2,455 | 1,342 | 1,342 | 1,321 | Cheng et al. (2017) |
| Nelson et al. | 3,073 | 3,073 | 1,378 | 735 | 671 | |
| Zhao et al. | 2,460 | 1,533 | 1,054 | 1,053 | 1,019 | Zhao et al. (2018) |
| PLNlncRbase | 51 | 51 | 33 | 33 | 31 | Xuan et al. (2015) |
| NONCODEv5 | 3,763 | 3,763 | 1,840 | 1,840 | 1,120 | Fang et al. (2018) |
| CANTATAdb | 4,373 | 4,373 | 1,083 | 1,083 | 1,011 | Szczesniak et al. (2016) |
| Liu et al. | 6,480 | 6,480 | 4,265 | 4,265 | 4,221 | Liu et al. (2012) |
| In-house data | 11,249 | 11,249 | 525 | 525 | 509 | |
| Total | 33,904 | 32,977 | 11,520 | 10,876 | 9,903 | |

**Supplemental Figure S1. Integration of lincRNA resources. (A)** Pipeline for the integration of lincRNAs from a variety of sources. **(B)** The table displays the number of transcripts that are remained at each step in our pipeline for each resource.

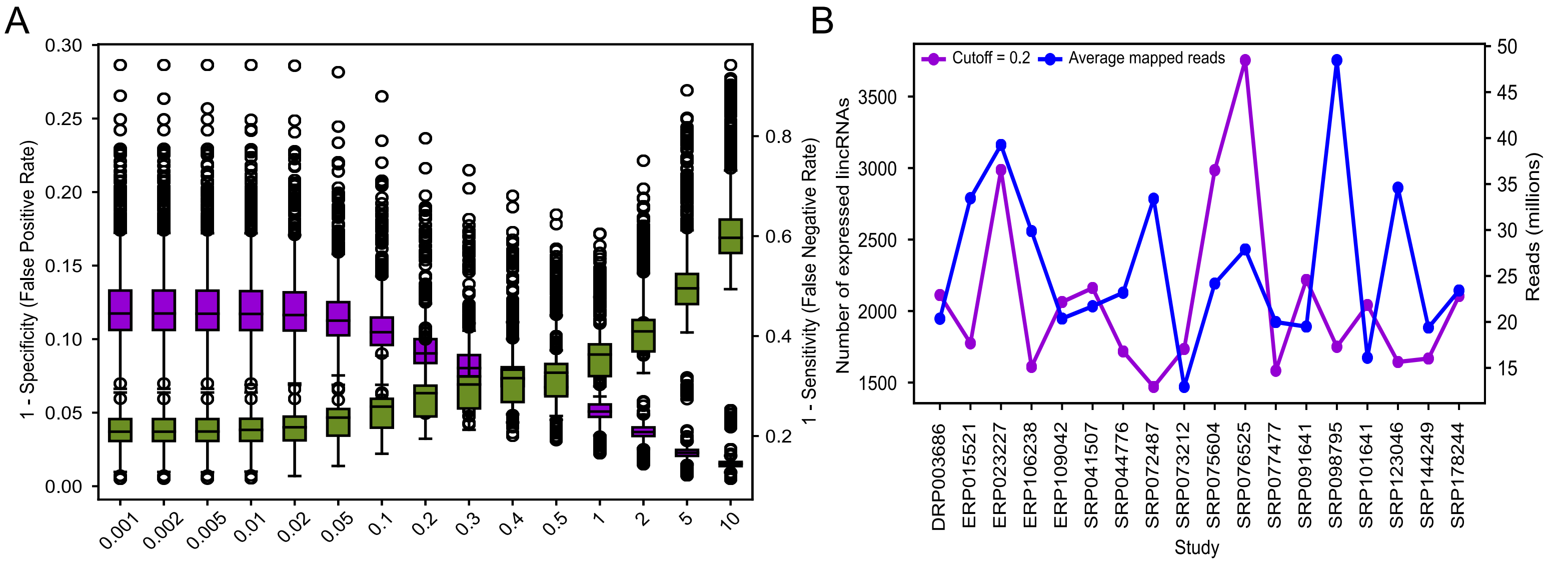


**Supplemental Figure S2. Definition of expressed genes and influence of sequencing depth. (A)** The protein-coding genes were considered as true positives, while the lincRNA genes were considered as true negatives. The false positive rate (purple) and false negative rate (green) were measured at different TPM thresholds for each sample. **(B)** The two lines show the distribution of number of the expressed lincRNAs (TPM Cutoff>0.2) (purple) and average number of mapped reads (blue) for per study.

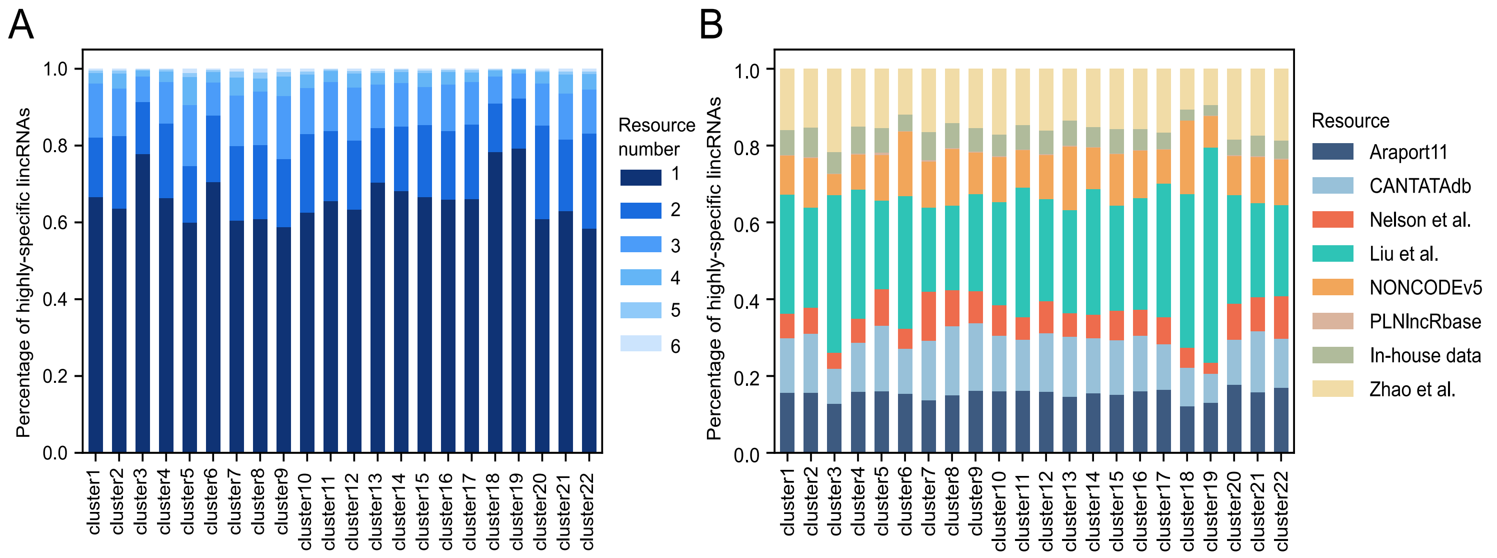


**Supplemental Figure S3. Resource annotation for highly-specific expressed lincRNAs. (A)** Percentage distribution figure showing the support for the number of resources for highly-specific lincRNAs in each cluster **(B)** Distribution of the percentage of highly-specific lincRNAs genes supported by each resource.

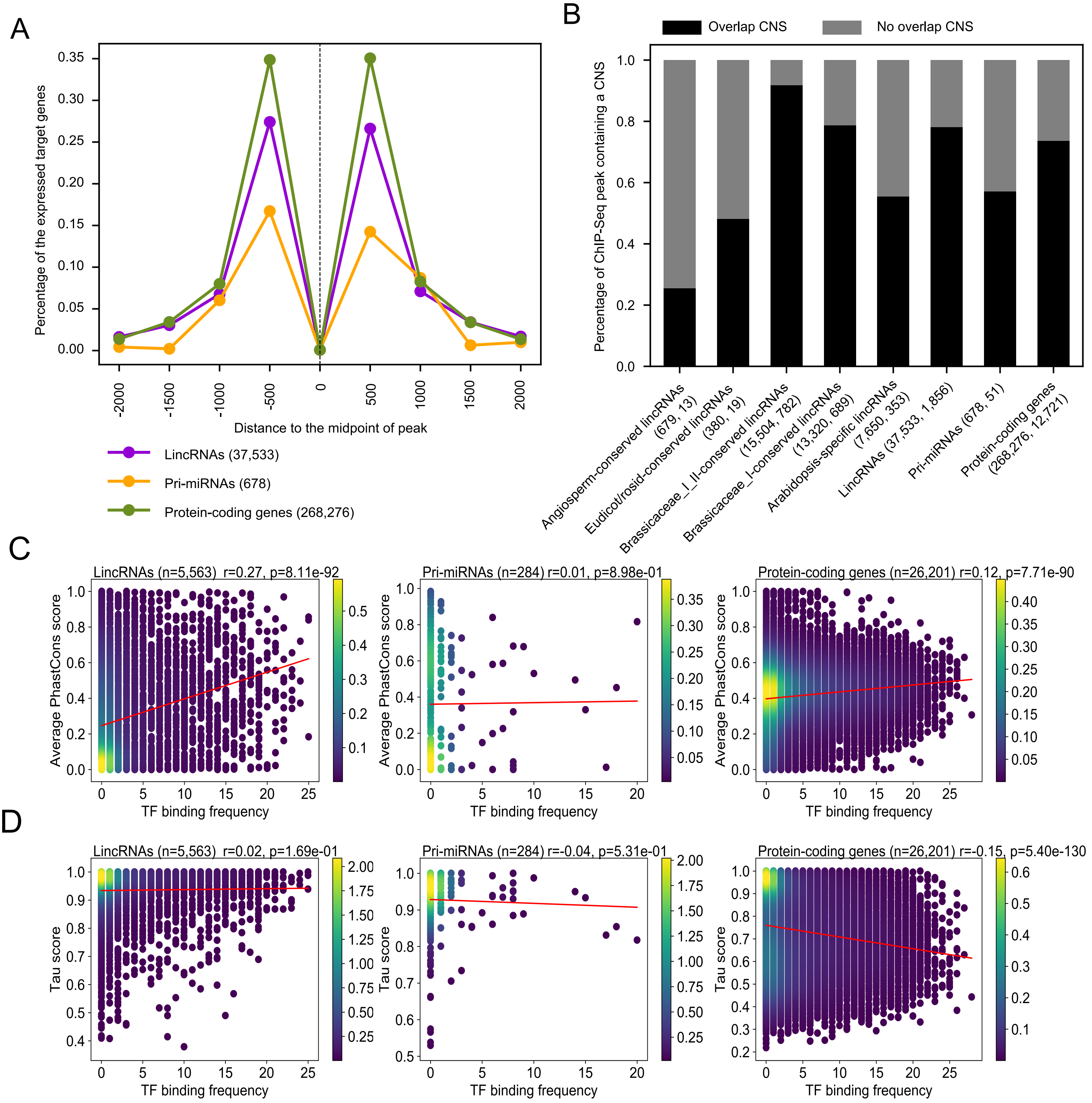


**Supplemental Figure S4. Regulatory properties of lincRNAs. (A)** Distribution of the distance of target genes from the peak midpoints for per gene type. The number in parentheses indicates total number of target genes <2kb from the peak midpoint. **(B)** Distribution of ChIP peak for per gene type overlapping with conserved noncoding sequence defined by Haudry et al. (2013). The first number in parentheses indicates the total number of the ChIP peak per gene type and the second number indicates the actual number per gene type containing at least one CNS. **(C-D)** Correlation of average PhastCons score and Tissue specificity (Tau) with TF binding frequency for lincRNAs, protein-coding genes and pri-miRNAs.

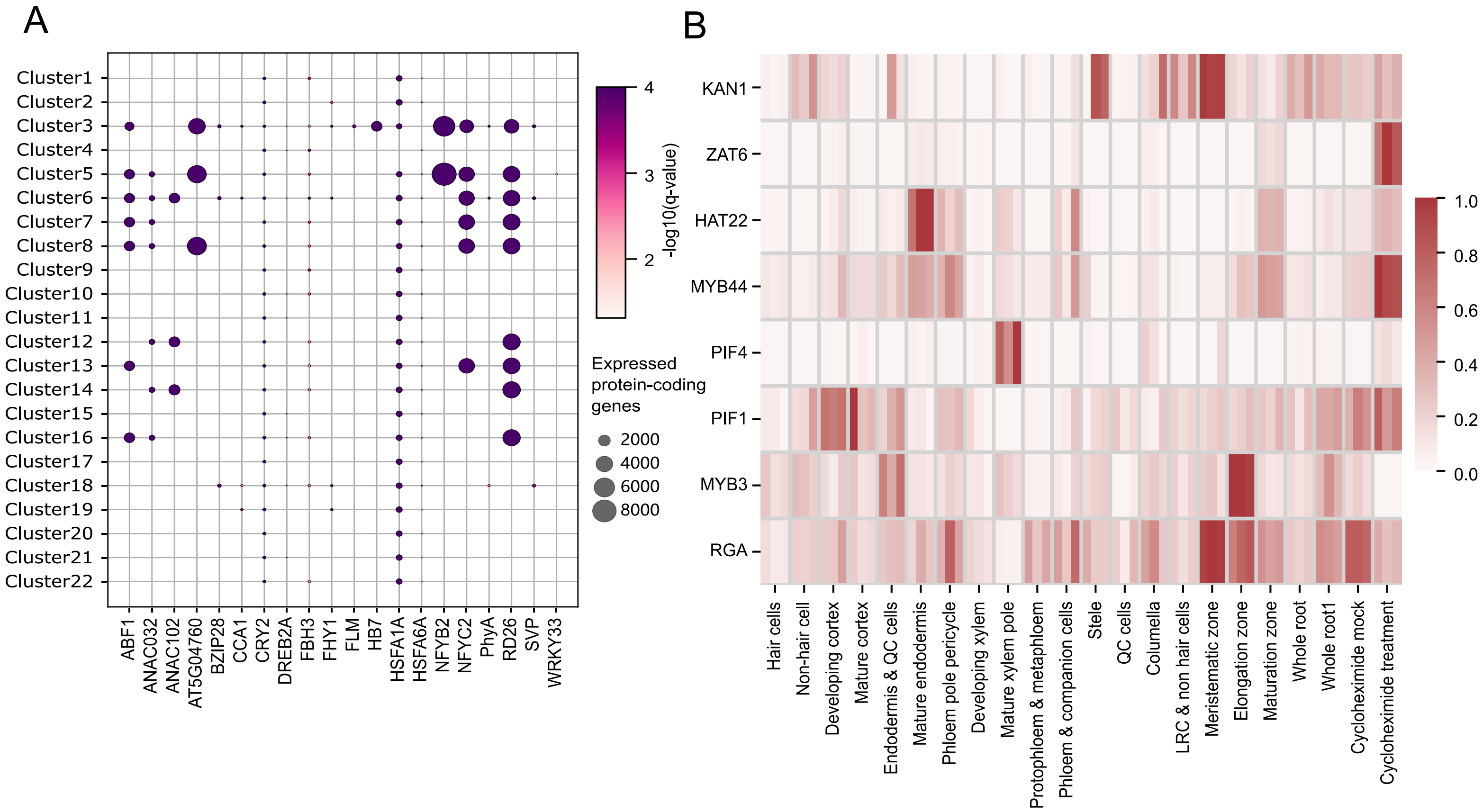


**Supplemental Figure S5. TF binding and expression. (A)** Bubble chart showing the enrichment of TF binding for expressed protein-coding genes in different expression clusters. TFs lacking significant enrichment in any of the 22 clusters are not shown. The dot sizes represent the number of the protein-coding genes while the color represents the statistical significance. **(B)** Heatmap showing TF expression in the cluster 19 root expression atlas. The color scale represents the normalized expression values (normalization means that the expression value in each row is divided by the row's maximum).

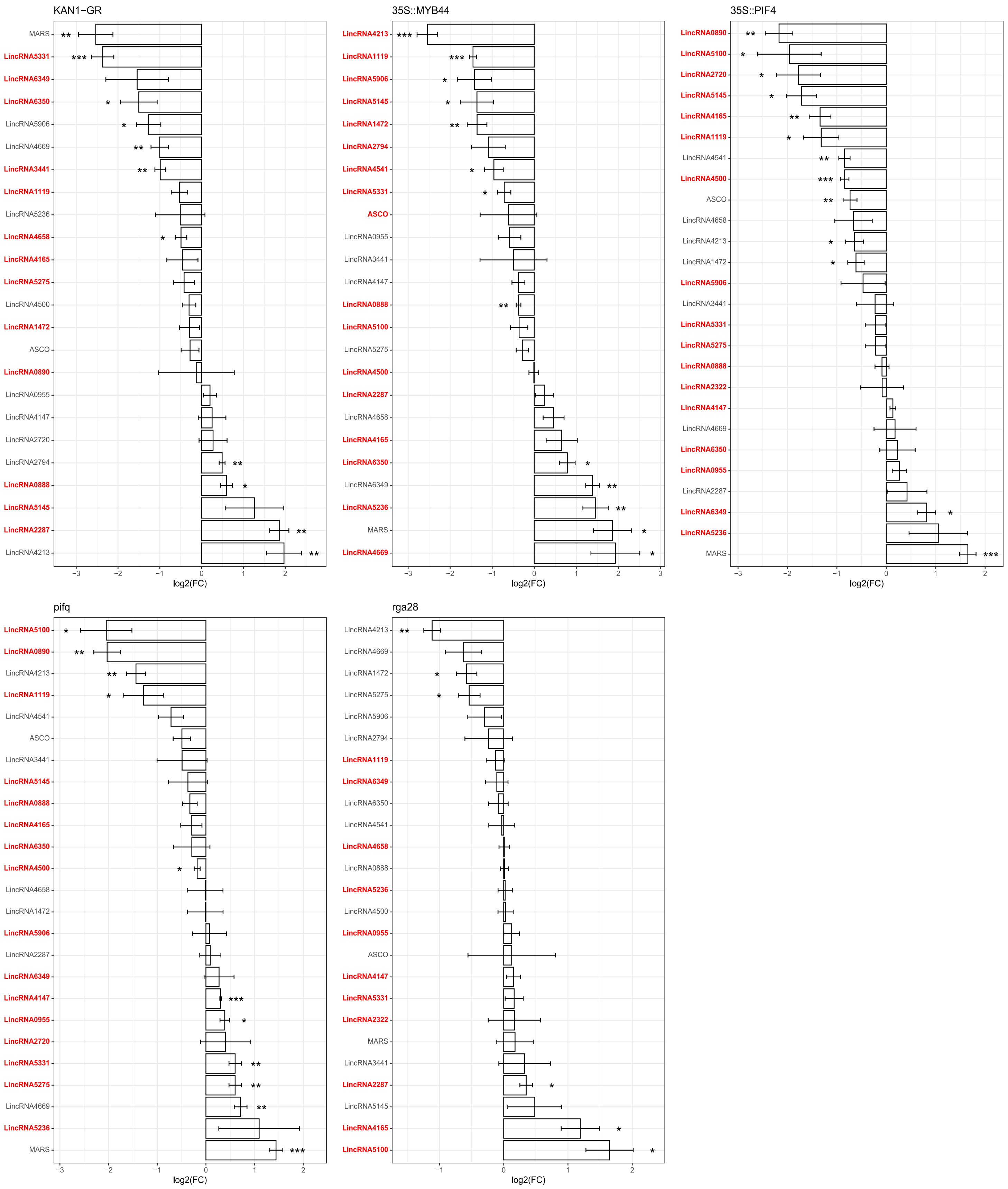


**Supplemental Figure S6. RT-qPCR validation results.** Bar charts showing log2 fold change (FC) of lincRNA relative expression levels in transcription factor (TF) overexpressing lines (35::*MYB44*, 35::*PIF4* and *KAN1-GR*) or TF knockout lines (*rga28* and *pifq* (*pif1/pif2/pif3/pif4* quadruple mutants)) versus control in 14-day old roots. Wild type lines were used as control. For the estradiol inducible *KAN1-GR* expression lines, DMSO treatment was used as control. Expression values were determined by RT-qPCR. PP2A was used as reference transcript for normalization. Asterisks indicate statistically significant differences (*p ≤ 0.05, **p ≤ 0.01, ***p ≤ 0.001) in an unpaired two-tailed Student's t-Test (n =3). Red color indicates TF ChIP peaks <2kb from the lincRNA gene. Conversely, grey color indicates lincRNA gene without TF peak that may have occurred due to distances > 2 kb or was identified in only one ChIP-Seq replicate, or where the lincRNA was not the closest gene to the TF peak.

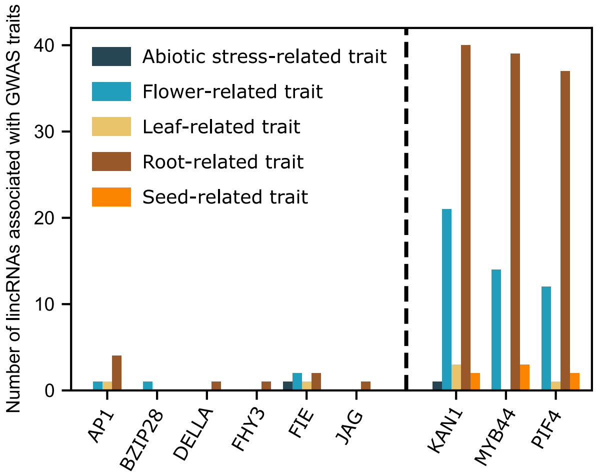


**Supplemental Figure S7. GWAS traits associated with lincRNAs confirmed by ChIP-Seq TF binding.** The bar plot displays the number of TF-regulated lincRNAs associated with each GWAS trait category. The lincRNAs regulated by the TFs on the right side of the black line show significant enrichment for TF binding in root cluster19, while the other TFs, shown on the left, lack enrichment for TF binding in root cluster 19.

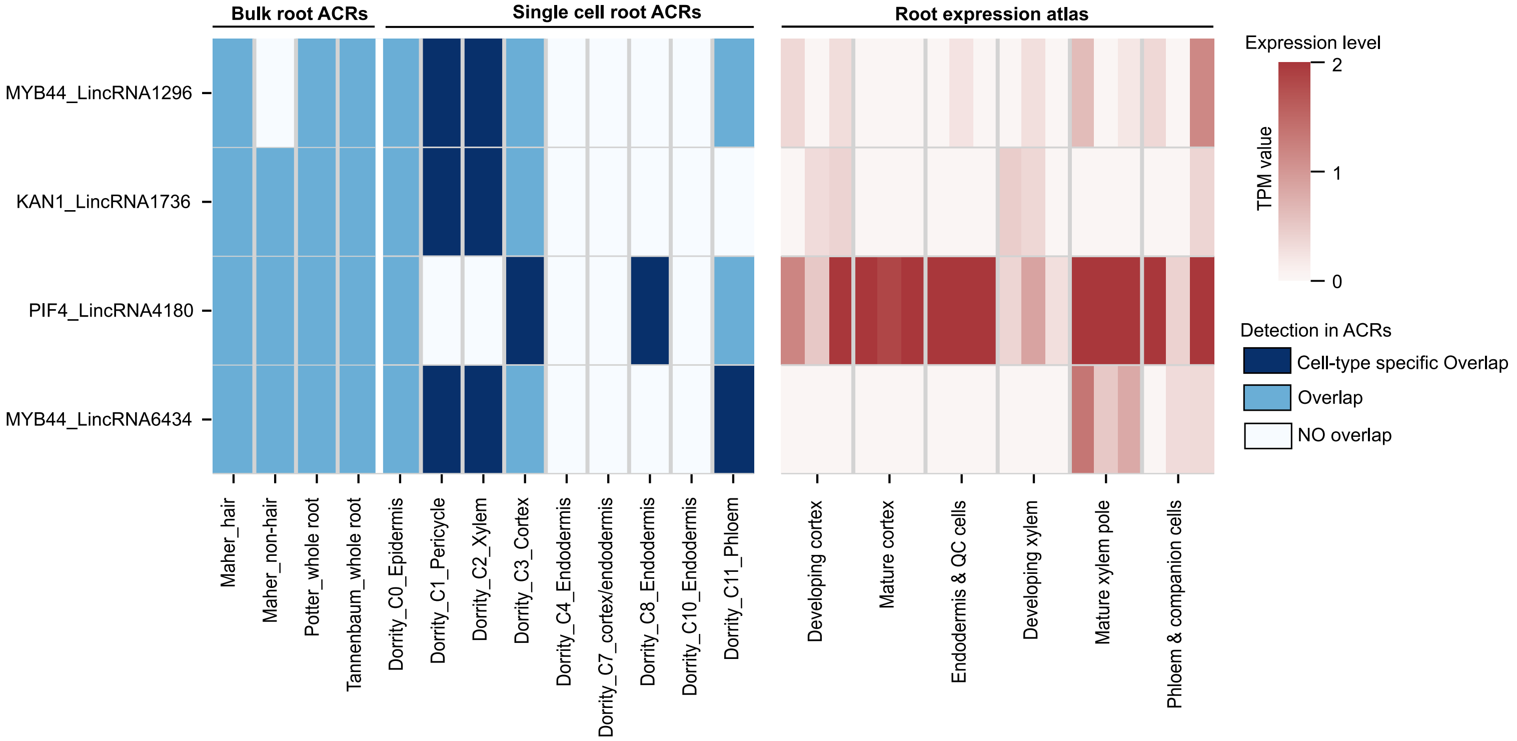


**Supplemental Figure S8. Identification of cell type-specific TF-lincRNA regulatory interactions.** Heatmap showing the lincRNA expression and ACRs confirm a regulatory interaction in the same cell type. The color scale represents the expression levels of lincRNAs in root cells. Three colors represent the detection of whether regulatory interactions overlap with ACRs (dark blue, blue, and light blue represent cell-type specific peak overlap, overlap, and no overlap, respectively).
